# Evolutionary history of New World monkeys revealed by molecular and fossil data

**DOI:** 10.1101/178111

**Authors:** Daniele Silvestro, Marcelo F. Tejedor, Martha L. Serrano-Serrano, Oriane Loiseau, Victor Rossier, Jonathan Rolland, Alexander Zizka, Alexandre Antonelli, Nicolas Salamin

## Abstract

New World monkeys (parvorder Platyrrhini) are one of the most diverse groups of primates, occupying today a wide range of ecosystems in the American tropics and exhibiting large variations in ecology, morphology, and behavior. Although the relationships among the almost 200 living species are relatively well understood, we lack robust estimates of the timing of origin, the ancestral morphology, and the evolution of the distribution of the clade. Here we integrate paleontological and molecular evidence to investigate the evolutionary dynamics of extinct and extant platyrrhines. We develop an analytical framework to infer ancestral states, the evolution of body mass, and changes in latitudinal ranges through time. Our results show that extant platyrrhines originated some 5–10 million years earlier than previously assumed, likely dating back to the Middle Eocene (∼ 43 million years ago, Ma). The estimated ancestral platyrrhine was strikingly small – weighing ∼ 0.4 kg, as compared to the largest modern species over 10 kg – matching the size of their presumed Eocene North African ancestors. Small-sized callitrichines (marmosets and tamarins) retained a small body mass throughout their evolutionary history, thus challenging the hypothesis of phyletic dwarfism as an explanation to their adaptive traits. In contrast, a rapid change in body mass range took place as the three families diverged between the Late Oligocene and the Early Miocene. That period also marks a peak in diversity of fossil platyrrhines and is associated with their widest latitudinal range, expanding as far to the South as Patagonia. This geographic expansion is temporally coincident with a significant increase in platyrrhine population size inferred from genomic data, and with warm and humid climatic conditions linked to the Miocene Climatic Optimum and the lower elevation of the Andes. These results unveil the early evolution of an iconic group of monkeys and showcase the advantages of integrating fossil and molecular data for estimating evolutionary rates and trends.

## Introduction

The platyrrhine primates, or New World Monkeys (NWM), are a diverse group of mammals currently distributed in the Neotropical region from Mexico to Northern Argentina but excluding the Caribbean Islands. They are all arboreal, but exhibit a wide spectrum of locomotor postures as well as body sizes (1). Most scholars accept that platyrrhines are divided into three families (2): Atelidae (including howler, wooly, spider, and wooly spider monkeys), Cebidae (including squirrel monkeys and capuchins, and marmosets and tamarins), and Pitheciidae (sakis and uakaries) (but see (3) for the discussion on the fourth family Callitrichidae).

Platyrrhines are thought to have originated in Africa, from which they dispersed into South America probably during the middle or late Eocene. The oldest fossil record, described as *Perupithecus,* was recently discovered in Santa Rosa, Peru, and is estimated to be of late Eocene age (4). The evidence for an African origin is supported by the exceptional morphological similarity between *Perupithecus* and the North African late Eocene *Talahpithecus* (4). These findings reinforce the hypothesis of a trans-Atlantic dispersal event during the Eocene, probably by rafting on budding forest islets (5, 6).

The fossil record of NWM is relatively diverse (33 extinct genera; (7)) but scarce in proportion to other mammals occurring in the same localities. In addition to *Perupithecus,* ancient records of platyrrhines also include Oligocene specimens from Contamana, Peru (8), and Salla, Bolivia, ca. 26 Ma (9-12). Patagonian and Chilean forms are known from the Miocene, ca. 20-15.8 Ma (7). These records conform to what is recognized as a first stage in platyrrhine evolution, with primitive and, in some cases, odd morphologies and often unclear phylogenetic positions (13). It was not until the Middle Miocene of Colombia, in the renowned fossiliferous area of La Venta, that the crown platyrrhines started to evolve into anatomically more modern forms, with morphologies in some cases indistinguishable from some living genera (7, 14).

Despite the fragmentary nature of NWM’s fossil record, available paleontological evidence hold the potential to reveal the evolutionary history of the clade. In particular, the geographic localities where platyrrhine fossils have been found indicate that past populations expanded into the Caribbean (Hispaniola, Cuba and Jamaica, where they no longer exist), and as far south as Patagonia, the southernmost area where non-human primates ever lived (12, 15). Other important insights about platyrrhine evolution can be obtained from the body mass of extinct taxa, which is strongly linked (and therefore predictable) from the molar area (16, 17). This allows a confident estimate of the body mass of extinct primates, even when the fossil record is extremely sparse. The fossil record of platyrrhines show that extinct taxa account for the largest (> 20 Kg) and some of the smallest (∼ 0.4 kg) taxa in the clade (as compared to the current range, approximately 0.1 to 12 kg), but the evolutionary dynamics shaping this wide range of body size changes remain unclear.

As an important complement to the fossil record, phylogenetic comparative methods based on molecular data and current trait measurements can shed further light into evolutionary processes (18). However, there are serious limitations to estimating ancestral states based on phenotypes and distributions of extant taxa only (19-22). In particular, for quantitative traits, ancestral states inferred from extant taxa cannot be estimated to be outside the observed range and tend be an average of the observed values, without the possibility to infer evolutionary trends (22-24). These issues should be particularly evident for NWMs, where the spectrum of body sizes and the geographic ranges are larger in extinct taxa than among living species.

Here we compile all available paleontological evidence and combine it with data from living species to infer the evolutionary history of NWMs. Based on a large molecular data set and available taxonomic information, we infer phylogenetic trees of extinct and extant lineages using the fossilized birth-death (FBD) method (25). We then analyze the history of two quantitative traits in the evolution of platyrrhines: body mass and the mean latitude of their geographic range. The abundance of teeth in the fossil record as compared to other skeletal parts, allows us to infer body mass for all described extinct taxa. The location of the fossil sites also provides valuable information on the geographical evolution of the clade. We develop a Bayesian framework to infer the evolutionary history of quantitative traits using phylogenies that incorporate both extant and extinct lineages, which we validate through extensive simulations. The method allows us to jointly estimate the ancestral states at each node in the tree and the rate (describing how fast a trait evolves) and trend (the tendency of a trait to evolve in an estimated direction, e.g., towards larger or smaller values). Both rate and trend parameters can change across lineages in the tree, and our algorithm jointly estimates the number and placements of shifts in parameter values.

By combining molecular and fossil evidence with our new method, we provide new insights into the origin and evolution of extinct and extant platyrrhines. Our analyses reveal how body mass evolution and changes in geographic distributions contributed to shaping the large diversity of life forms encountered today in New World monkeys.

## Results

### NWM phylogeny

Our phylogenetic analysis of the platyrrhine clade encompassed 87 extant species and 34 extinct taxa, spanning from the Late Eocene to the Holocene (Tables S4-5). The phylogenetic relationships between extant species were highly supported (Figs. S6-7) and reflected the results of Springer*, et al.* (26), while the placement of extinct taxa was sampled by the FBD within the limit of taxonomic constraints (see *Methods*). The FBD analysis placed the origin of the clade in the Eocene and the divergence times of families and subfamilies between the late Oligocene and the mid-Miocene (Table 1). The speciation and extinction rates estimated as part of the FBD model indicate a high turnover rate (extinction / speciation rates = 0.75, 95% credible interval: 0.57–0.91) which is consistent with the relatively high diversity of extinct taxa (Table S6).

**Table 1.** Estimated branching times summarized for the main clades within platyrrhines. The origin of the clade corresponds to the stem age of Platyrrhini as estimated by the fossilized birth-death model.

| **clade** | **Age (Ma)** | **95% HPD** |
| --- | --- | --- |
| Origin of the clade | 43.53 | (37.64-50.77) |
| Crown age of Platyrrhines | 32.84 | (27.43-38.60) |
| Crown age of Pitheciidae | 25.11 | (21.01-28.88) |
| Crown age of Cebidae | 24.05 | (20.98-27.29) |
| Crown age of Cebinae | 22.33 | (20.93-25.00) |
| Crown age of Pitheciinae | 21.76 | (20.00-25.10) |
| Crown age of Aotinae | 20.91 | (20.00-23.69) |
| Crown age of Homunculinae | 19.49 | (17.00-23.76) |
| Crown age of Atelidae | 18.36 | (14.26-22.58) |
| Crown age of Callitrichinae | 17.60 | (14.38-20.82) |
| Crown age of Alouattinae | 15.09 | (12.50-18.29) |
| Crown age of Atelinae | 11.99 | (8.96-15.47) |

### Methodological validation

Extensive simulations show that our novel Bayesian method to infer trait evolution along a phylogeny with extinct and extant taxa provides accurate estimates of the rate and trends parameters and their variation across clades (*Supplementary Information*). Our algorithm correctly identified the number of shifts in rate or trend (if any) with a frequency of 91.5% (ranging from 79 and 98% across different simulation settings; Table S1). The mean absolute percentage error (MAPE, see Methods) ranged from 0.11 to 0.22 for the rate parameter and between 0.11 and 0.23 for the trend parameter, in data sets with 20 fossils included (Fig. S2a,b, S4d,e). The estimated ancestral states were very accurately estimated with average coefficients of determination (R^2^) ranging from 0.92 to 0.99 across simulations (Fig. S2c, S4c). Decreasing the number of fossils included in the data had small effects on the accuracy of the estimated rate and trend parameters (Fig. S3a,b) and on the ancestral states (Fig. S3c). In the absence of fossil data, the method could still estimate accurately the rate parameter (while the trend parameter is unidentifiable), but the accuracy of the ancestral states decreased to R^2^ = 0.82 for data sets simulated without trends, and to R^2^ = 0.53 for traits evolving under a non-zero trend (Fig. S2). The performance of our algorithm in terms of efficiency of sampling the parameters from their posterior distribution and time to evaluate the likelihood significantly outperformed alternative implementations, reducing computation times by one order of magnitude (Fig. S5). Notably, whereas computation time of alternative implementations increases exponentially with tree size, it increases linearly with our method, thus allowing for efficient analysis of very large data sets (thousands of tips; Fig. S5).

### Body mass

Platyrrhine body mass was estimated to have evolved under a Brownian motion (BM) model with variable rates and no or little evidence for positive or negative trends. The estimated number of rate shifts varied among the 100 FBD phylogenies analyzed and ranged between 0 and 7, whereas the trend parameter was found to be mostly homogeneous across branches (Table S7). We found comparatively high rates of body mass evolution at the family level, whereas the rates were substantially lower within subfamilies, except for Cebinae, for which a high rate was inferred (Fig. S8). The trend parameters were overall very close to 0, indicating non-directional evolution of body mass. The ancestral body mass inferred at the root of the tree ranged between 260 and 890 g (Fig. 1; Table S8). This estimate strongly differs from the estimate obtained from a phylogeny of extant NWMs, after discarding the fossil record (Fig. S10), where the estimated ancestral body mass at the root ranged between 949 and 2,710 g (Table S8).

**Figure 1.**
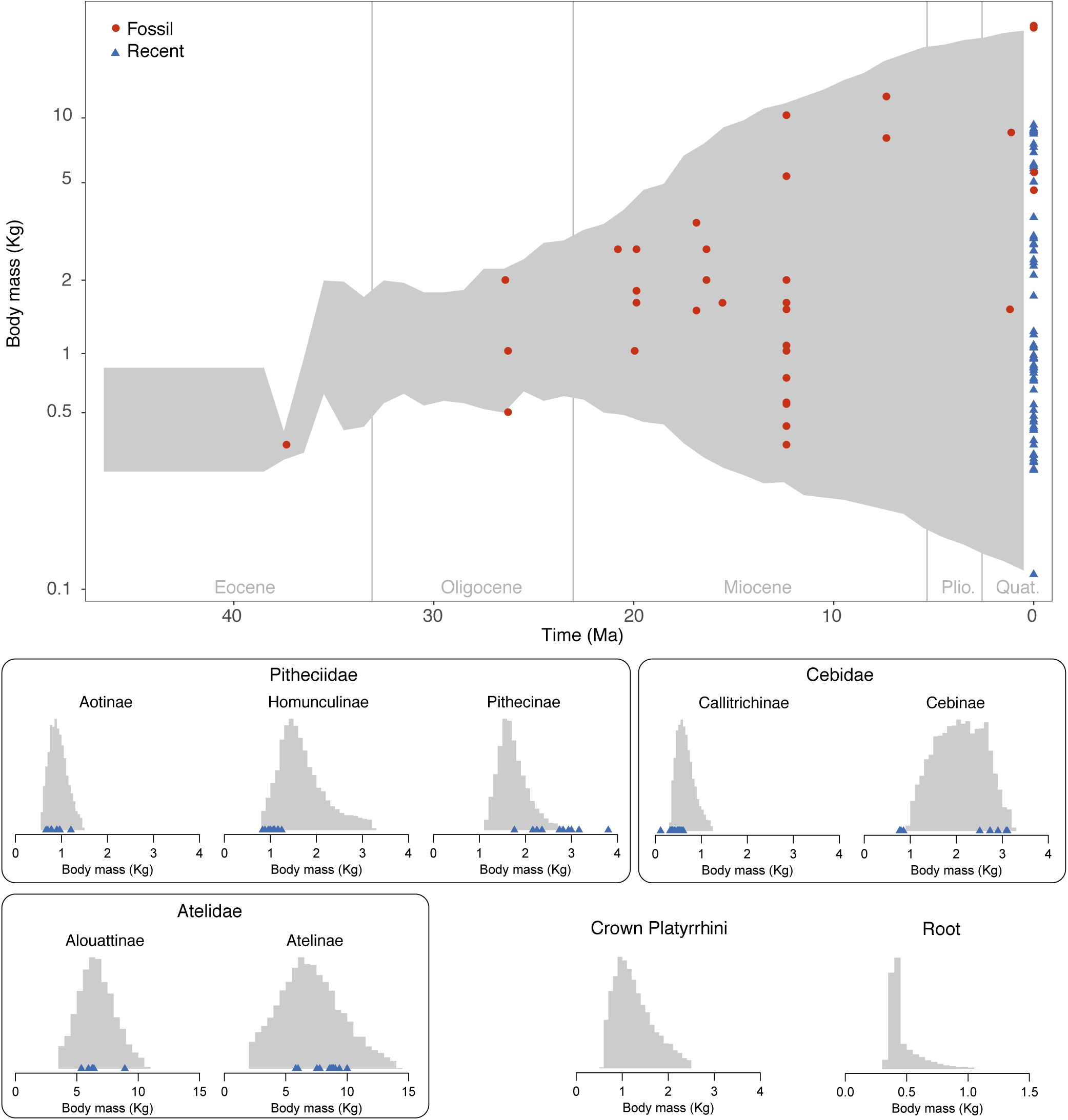
Body mass evolution in platyrrhines. The gray shaded area in panel A) shows the range of trait values through time (95% credible interval) inferred across a sample of 100 phylogenies with extinct and extant taxa. The red circles indicate the log body mass and age of the fossil taxa included in the analysis, and the blue triangles show the log body mass of extant species. The histograms in panel B) show the estimated ancestral body mass (posterior distributions truncated to their 95% credible intervals) across subfamilies, for the most recent common ancestor of extant platyrrhines and at the root of the tree.

### Range occupancy

Latitudinal ranges in platyrrhines have changed at variable rates across lineages and we estimated 1 to 4 rate shifts among the 100 FBD phylogenies analyzed (Table S7). The rates were highest in the subfamilies Alouattinae and Cebinae (Figure S9). The trend parameter was inferred to be constant across clades and positive, although it did not differ significantly from 0 based on the 95% CI (Table S7, Figure S9). The inferred ancestral latitudes through time show an expansion to the South which culminates in the Early Miocene. The slightly positive trend likely captures the general trend of ancestral latitudes towards the present tropics following the disappearance of platyrrhines from the south of South America around the Middle Miocene and expansion into Central America during the Miocene and Pliocene (Fig. 2, S9).

**Figure 2.**
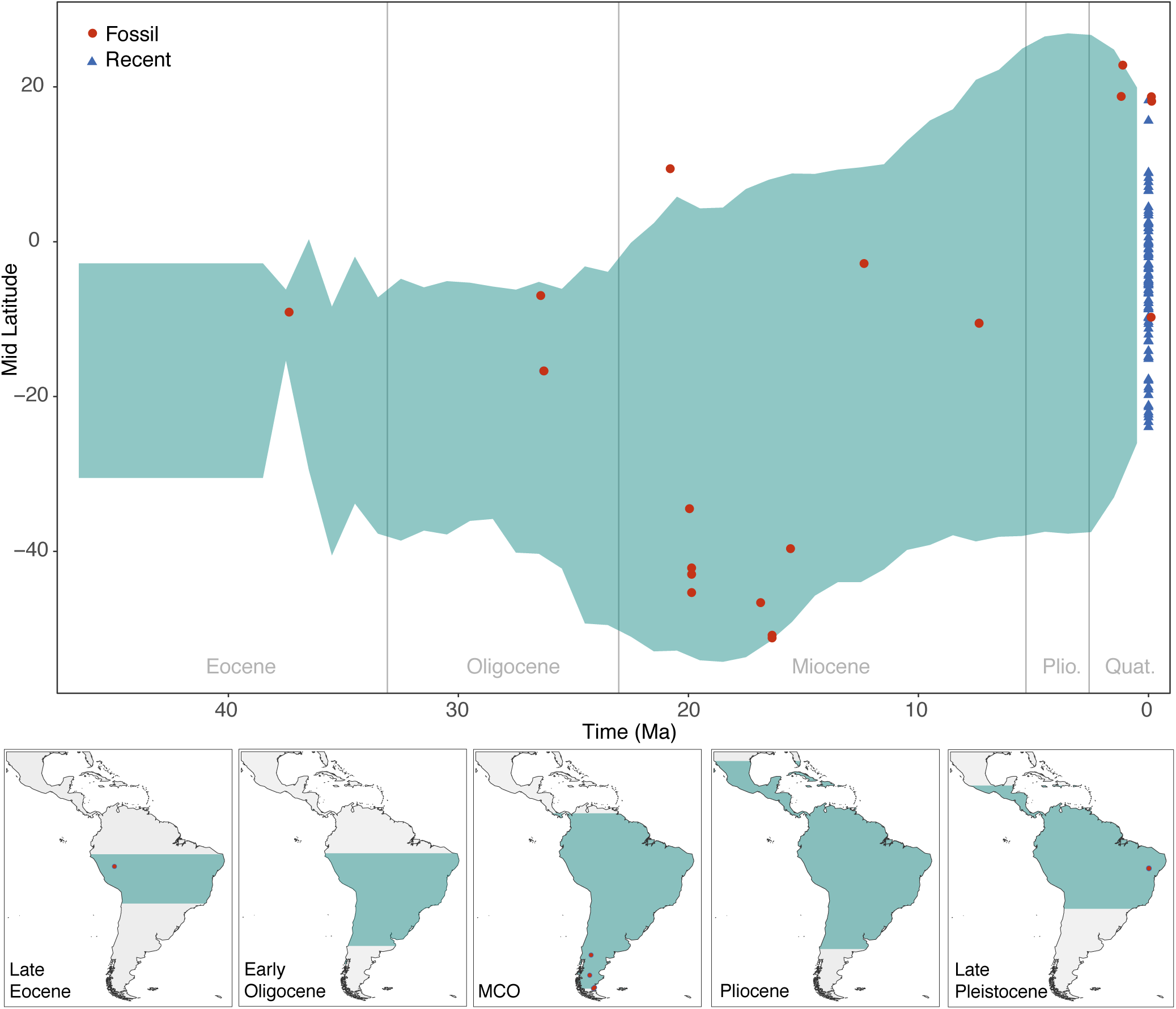
Changes in latitudinal ranges. The gray shaded area in panel A) shows the range of trait values through time (95% credible interval) inferred across a sample of 100 phylogenies with extinct and extant taxa. The red circles indicate the (present day) latitude of the localities of the fossil taxa included in the analysis, and the blue triangles show the mid latitude of the geographic range of extant species. The histograms in panel B) show the estimated mid latitudes (posterior distributions truncated to their 95% credible intervals) for major nodes of the tree.

## Discussion

### Origin and body size evolution

Our phylogenetic analysis of the platyrrhine clade provided generally older estimates of divergence times compared to previous studies (12, 26-28). For instance, the crown age of all extant platyrrhines was pushed back from the Early Miocene of previous estimates to the Early Oligocene. This is likely the result of using a more substantial and complete amount of fossil information and a more realistic approach to calibrate the tree (25, 29). The estimated time of origin of the platyrrhine lineage (stem age) is 43 Ma, thus indicating that the morphologically similar extinct North African anthropoids are comparable in age (30)(Table S10). This finding suggests that platyrrhines (or pre-platyrrhines) evolved first in Africa, where they eventually went extinct, and dispersed sometime in the Eocene into South America where they diversified, in the absence of other primates in the continent (4). Previous estimations also suggested an Eocene age for the catarrhine-platyrrhine split (2, 27). Based on these lines of evidence, it is probable that the NWMs origin (stem age) dates back to at least the Middle Eocene.

Ancestral state estimates indicate that NWMs derive from a remarkably small ancestor (around 400 g; Fig. 1), with an inferred body mass close to the lower boundary of the size range of extant platyrrhines. In addition to *Perupithecus*, the oldest fossil included in this study, fossil records from the same locality (Santa Rosa, Peru) include two broken upper molars and a lower molar of different unidentified taxa (4). Their damaged condition did not allow an accurate description of these taxa and the specimens were therefore not included in our analyses. However, their approximate body size was likely around 70% of the size of *Perupithecus*. Thus, the taxon was possibly the size of a living tamarin, such as *Callimico* or *Saguinus*, weighing about 280 g. Despite the uncertainties around these estimates, this fragmentary fossil evidence indicates that the body mass of the ancestral NWM might have been even smaller than 400 g and closer to the size of small marmosets *Mico*, *Callithrix* and *Cebuella*. The hypothesis of a small ancestral body mass at the origin of NWMs is also supported by the estimated size of Eocene anthropoids, especially parapithecoids from North Africa, which are mostly around 500 g (1, 31), with *Talahpithecus* weighing less than 400 g (Table S10). We propose that a small body size was probably a key factor enabling the survival of the individuals that reached South America from Africa, since their resource requirements on a floating islet would have been substantially smaller (e.g., relying on a diet of invertebrates and being better capable of protecting themselves against dehydration) than for a heavier and larger organism.

NWMs reached larger body sizes above 2 kg between the Late Oligocene and Early Miocene. Then, the upper bound of the body mass range in NWMs continuously raised, eventually reaching now extinct giant forms in the Atelidae family like *Cartelles* or *Caipora*, from the Pleistocene of Brazil (> 20 Kg; Fig. 1; Table S5). Despite the expansion in range of platyrrhine body mass through time, most clades maintained intermediate sizes around 1 – 3 Kg (Fig. 1). Most known fossil species until the Miocene, including all the taxa found in Patagonia displayed intermediate body mass (e.g. pitheciids *Homunculus* and *Carlocebus* and cebins *Dolichocebus*, *Killikaike*; Table S5). Body size evolution in platyrrhines was largely heterogeneous, as indicated by several estimated rate shifts (Table S7), probably as a response to differential evolutionary selection forces acting on species across the vast range of Neotropical habitats.

Asides from the Late Miocene cebid *Acrecebus fraileyi* (12 Kg), most large NWMs belong in the family Atelidae. The large body size of atelids is associated with particular locomotor adaptations, some of them displaying suspensory behavior, although the four living genera have prehensile tails (1). Living atelids are widely distributed from Central America through northern Argentina, and some taxa can cope with open environments and seasonal climates, such as *Alouatta*. Despite the wide range of ecologies and adaptations in modern atelids and their large geographic range, the fossil record of the family does not include any specimen from the Miocene sites in Patagonia suggesting that they did not expand as far south as cebids and pitheciids (but see 32).

The smallest NWMs, the callitrichines (marmosets and tamarins), are characterized by a body mass smaller than 1 kg, a distribution spanning tropical forests from the Amazonia to Central America, and very scarce fossil record. The small body size of callitrichine has been usually considered as the result of evolutionary dwarfism (Ford, 1980, 1986), and this evolutionary trend is assumed to explain their atypical morphology, such as the reduction or loss of the third molars, loss of the hypocone, presence of claws instead of nails. However, in our analyses, we did not find evidence of a negative trend in body mass evolution within the subfamily (Fig. S8). Furthermore, the estimated size of the common ancestor of living callitrichines is small (around 0.6 kg; Fig. 1, Table S8), suggesting that callitrichines may have maintained a small body size throughout their evolutionary history. These results challenge the hypothesized dwarfism in callitrichines, and suggest that their peculiar morphological features might be the result of ecological adaptations, which did not necessarily involve a reduction in body mass.

The process of body size evolution was heterogeneous across platyrrhine clades and did not follow a simple neutral evolution. We found evidence for several rate-shifts but only weak or negligible trends. Because we incorporated topological uncertainties in our analyses, which is quite substantial for extinct lineages, it is difficult to pinpoint the exact placement of rate changes. However, we do observe that the rate of body mass evolution is highest at the family level and lower within most subfamilies (Fig. S8). This suggests a rapid change in body mass range as the three families diverged, between the Late Oligocene and the Early Miocene, followed by a slower pace in evolution within subfamilies (Fig. S8).

### Diversity dynamics and geographical occupancy

Our analyses show that the Late Oligocene and Early Miocene also mark an important phase in the biogeography of NWMs, with a peak in fossil diversity and the widest latitudinal range in South America. The geographic expansion of platyrrhines started in the Late Oligocene and culminated between 20 and 15 Ma, when they reached their southernmost distribution. This geographic expansion is temporally coincident with a significant increase in NWMs population size as inferred in a recent analysis of genomic data (2). Thus, both fossil and molecular data identify this phase as a crucial event in platyrrhine diversification and evolution.

During the Early Oligocene, the Patagonian mammal fauna had experienced a marked turnover (the “Patagonian Hinge”; (33)) associated with global climate cooling, during which Polydolopimorformes, a diverse group of marsupials widely distributed in Patagonia and across South America, declined and eventually went extinct during the Early to Late Oligocene (33, 34). Among the Polydolopiformes, several groups showed evident primate-like adaptations in diet and probably paleoecology, and their extinction might have played a role facilitating the colonization of primates across Patagonia prior to the Early Miocene.

Climate change likely had a direct effect in determining NWMs changes in latitudinal range. Modern platyrrhines are mostly adapted to tropical environments and their occurrence in Patagonia around 20–16 Ma was likely allowed by the warmer climate characterizing the Miocene Climatic Optimum (MCO) (35). Additionally, this period precedes a phase of major Andean uplift and the lack of a high elevation mountain range likely meant more humid paleoenvironments in the region, with moisture coming from the Pacific Ocean (36). The MCO was followed by the diastrophic Quechua phase (37), in which the regional climate and environment in the south of South America ware affected both by global cooling and by the onset of Andean uplift. The latter progressively interrupted the influx of moisture from the Pacific and produced a rain shadow effect, thus turning Patagonia into a more open environment, with extensive grassland (38, 39) and causing a faunal turnover (13, 40, 41). The environmental change that took place in Patagonia through the Middle and Late Miocene coincides with a range contraction of the southern limit of platyrrhine distribution (Fig. 2). The absence of latitudinal barriers (e.g. mountain ranges, deserts) in the continent has likely contributed to making these range expansions and contractions happen (42). Although richer fossil occurrence data would be necessary to assess the roles of extinction and migration in this process, our estimates of high extinction rates in platyrrhines (Table S6) suggest that several Patagonian lineages have gone extinct after the MCO (12). This scenario allows us to establish analogies between the Miocene primates from Patagonia, and the southernmost living platyrrhine that reaches the northeast of Argentina today: the black and golden howler monkey, *Alouatta caraya*. This living primate inhabits open environments and sometimes patches of forests inside extensive grasslands affected by strong seasonality, including several freezing days during winter. In general, platyrrhines may be considered as indicators of tropical or subtropical environments, but the exception of *A. caraya* living in the most extreme conditions for a Neotropical primate, indicates that the Patagonian forms may have experienced similar climatic and environmental conditions.

During the Miocene and Pliocene we infer a range expansion of platyrrhines towards the north of South America and into Central America and the Caribbean, although the scarcity of primate fossil records limits our ability to assess the detailed dynamics of this range expansion (12). Nevertheless, our analyses, which included the recently described oldest record of NWM in North America (Tables S4-5) (28), support the hypothesis of a Miocene colonization of the North American tropics coincident with a complex and progressive closure of the Panama Isthmus (Fig. 2) (28, 43, 44).

### Methodological advances

Recent studies have shown that the inclusion of fossils in phylogenetic analyses of trait evolution can improve dramatically our ability to infer ancestral states and the underlying evolutionary processes (45). Here we implemented a powerful Bayesian analytical framework to integrate fossil and phylogenetic data in comparative analysis. Using this method, we showed by simulations that the estimation of ancestral states in the presence of evolutionary trends is far from realistic unless at least some fossil information is provided (Fig. S3).

Previous analyses of trait evolution combining data from extinct and extant taxa were either based on node calibrations (21) or on phylogenetic hypotheses built from morphological data (46, 47). Node calibrations are based on the assumption that a fossil can be confidently placed at a node in the phylogeny of the living descendants, i.e. it is not a stem or a side branch. The topological placement of fossils in a phylogeny (whether as nodal constraints or extinct tips) is difficult to determine in many clades especially when there is scarcity of morphological traits that can be scored from fossils, as in NWMs, or morphological traits bear little phylogenetic value (12, 48). In our study, we used the available taxonomic information to constrain the position of fossil lineages and, at the same time, relied on the FBD process to sample multiple phylogenetic hypotheses based on an explicit process of speciation, extinction and preservation. Thus, by running trait-evolution analyses on FBD trees we incorporated topological and temporal uncertainties in our estimates.

There remain, however, some limitations in inferring evolutionary history of traits based on the fossil record. The small number and fragmentary nature of available fossil occurrences make it difficult to appreciate the amount of intraspecific variability (e.g. differences in body size across populations or linked to sexual dimorphism) and the impact of measurement error, both of which may affect the reliability of the estimates (49, 50). In the case of biogeographic inferences, while more realistic and spatial-explicit models (e.g. 51) would be desirable to infer the biogeographic history of a clade, the lack of extensive occurrence records means that the analyses must rely on unavoidable simplifications and assumptions. For instance, most platyrrhine fossil taxa are known from single localities and we used these coordinates as representative of their latitudinal range. Despite the limitations associated with the use of fossils in comparative analyses, paleontological data are still crucial evidence of traits that no longer exist in the living descendants (21).

Furthermore, simulations using our Bayesian framework to infer trait evolution show that even very few fossils can drastically improve the estimation of ancestral states, indicating that the method can be applied to a large number of empirical data sets (Fig. S3). The importance of fossil information in the inference of evolutionary processes is evident in our analyses of body size and mean latitudinal range in platyrrhines, where the inclusion of fossil data significantly changes the estimated processes and ancestral states (Figs. S10-11, Tables S8-9).

## Conclusions

Our integrated analysis of molecular and fossil data of NWMs revealed that the stem platyrrhine lineage originated earlier than previously thought, and almost certainly in tropical regions. NWMs diversified shortly after their arrival from Africa and expanded their latitudinal range into the entire South American continent. This expansion was probably triggered by a global warming in the Miocene, which increased the extent of environmentally suitable regions for the diversification of mostly megathermal species such as monkeys. Later, global cooling contributed to the disappearance of NWMs from the southernmost part of South America, resulting in a contraction of the range around the modern tropics. Some primate populations living in seasonal environments are still distributed in northern Argentina, in open forests that may be analogous to the Early Miocene Patagonian environments. The remarkable (paleo)environmental diversity of Central and South America ranging from tropical forests to the Andean highlands through the more seasonal Patagonia, favored the remarkable body size diversification of the NWM, occupying different niches that avoided competition, and taking advantages of many resources for non-overlapping dietary adaptations.

Under the new evidences about the earliest ancestors of NWM, we infer that body mass evolution in these primates started with small forms, as shown by their Eocene North African relatives, but were later diversified in a wide range of large and middle-sized taxa. In contrast, callitrichines kept small sizes throughout their evolutionary history, challenging the widely-accepted hypothesis of phyletic dwarfism in this clade.

## Methods

### Bayesian analysis of trait evolution

We implemented a Bayesian algorithm to estimate the evolutionary history of quantitative traits in a phylogenetic framework. The evolutionary models implemented here are based on Brownian motion (BM) in which the expected trait value *v*_*i+t*_ at time *t* follows the normal distribution:

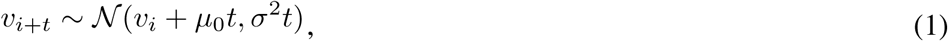

where *v*_*i*_ is the ancestral trait value at time *i*, *μ*_*0*_ is the trend parameter describing the tendency of a trait to evolve in a direction, and σ^2^ is the rate parameter describing the speed of phenotypic change. Note that most commonly neutral BM models are applied in the absence of fossil data by setting *μ*_*0*_ = 0. In our implementation, we relax the assumption of a constant BM model by allowing both the rate and the trend parameters to vary across clades in the phylogeny. We use a Birth-Death Markov Chain Monte Carlo algorithm (BDMCMC; 52, 53) to estimate the number of rate and trend shifts and their placement in the tree from the data (see *Supplementary Methods*). In addition to the BM parameters, our approach jointly estimates the ancestral states of the quantitative trait for all internal nodes. The likelihood of a vector of ancestral states ***v*** = [*v*_*1*_*, …v*_*N-1*_] (where *N* is the number of extinct and extant tips in the tree) is calculated as a product of normal densities based on Equation 1 and on the current values of ancestral states and BM parameters, recursively from the tips to the root (23). We used arbitrary vague normal priors on the ancestral states (with mean = 0 and standard deviation = 100), an exponential prior on the BM rates, and a normal distribution (with mean = 0 and standard deviation = 10) on the trend parameters. In order to estimate efficiently the ancestral states, we implemented a Gibbs algorithm to sample them directly from their posterior distribution (see *Supplementary Methods*), whereas a Metropolis-Hastings sampler was used to estimate the BM parameters and the root state (54). We thoroughly tested our implementation through extensive simulations (see *Supplementary Methods* and *Supplementary Results*) assessing 1) the robustness of model selection using BDMCMC, 2) the accuracy of parameter estimation, and 3) the performance of our algorithm compared to alternative implementations.

### Phylogenetic analysis

We used molecular dataset from Springer et al. (26) from which we kept only the 87 species of platyrrhines and discarded all markers with a high proportion of missing data for these taxa. The reduced alignment included 54 nuclear genes and 1 mitochondrial gene for a total length of 36,065 bp, with average coverage per gene of 60% of the living taxa (Tables S2-3). We used 34 fossil taxa with ages ranging from the Late Eocene to the Pleistocene (Tables S4-5) to infer a phylogeny of living and extinct platyrrhines. Phylogenetic relationships and divergence times were jointly estimated in BEAST v2.4.3 (55) under the Fossilized Birth Death model implemented in the Sample Ancestor package (25, 56). We selected a log-normal relaxed clock and used the same gene partitions as in Springer et al. (26) with GTR+Γ substitution models for each gene. Under the FBD model, fossil taxa can be treated as direct ancestors or extinct tips and their topological placement is treated as nuisance and integrated out using MCMC (25). We use taxonomic information following (7, 13) to constrain the placement of extinct taxa in the phylogeny when possible (e.g. to a family or subfamily; Table S4). We ran two MCMC analyses for 100 million generations, sampling every 10,000 generations. Both runs were examined in Tracer v1.5 to check for convergence. We combined the 2 runs after removing a burn-in of 25 million generations. To obtained a dated phylogeny with only extant species, we removed fossils taxa from the entire posterior distribution of trees and used them produce a maximum clade credibility tree.

### Trait analyses

We compiled fossil data and body mass estimates for most extant and extinct taxa from (1). We additionally obtained data for *Perupithecus* from (4), for *Canaanimico* from (8), for *Talahpithecus* from (31), and for *Panamacebus* from (28). Mean latitudes of extant species were computed from the minimum and maximum latitudes of their ranges as defined in the IUCN database (http://www.iucnredlist.org) and from the Pantheria database (57). Because most fossil taxa are known from single localities we used the latitude or their sampling locality as representative of their mean latitudinal range. We treated latitude as a quantitative trait to infer temporal changes in the ancestral distribution in platyrrhine evolution (e.g. 58).

We ran trait evolution analyses on 100 trees randomly selected from the BEAST posterior sample in order to incorporate topological and temporal uncertainties in our estimates. On each tree, we ran 5 million BDMCMC iterations sampling every 5,000.

Because the trees differed in branching times and in topology, we summarized the ancestral states for each family and subfamily (Table S9). We also calculated the range of trait values occupied through time as the minimum and maximum boundaries of the range of estimated ancestral states averaged over 100 analyses within 1 million-year time bins, following the procedure of (59). The number and placement of rate and trend shifts estimated through BDMCMC varied across trees. Thus, we summarized the parameters by families and subfamilies, by averaging the estimated rates and trends across the lineages within (sub)families. For comparison, we repeated the analysis of body mass and mean latitude evolution after dropping all extinct taxa from the platyrrhine phylogeny (Figs S10-11).

## Data and software availability

All the data used in this study (including the nucleotide alignment, BEAST input file and trait data) are available here: https://github.com/dsilvestro/fossilBM. The repository also includes all the R and Python scripts developed to simulate data and analyze them.

## Author contributions

DS, MT, and NS conceived the study, DS, MLS, AZ, OL, VR, JR, NS contributed to coding and analysis, DS, MT, AA, and NS led the writing.

## Acknowledgments

All analyses were run at the High-performance Computing Center (Vital-IT) from the Swiss Institute of Bioinformatics. D.S. received funding from the Swedish Research Council (2015-04748). M.F.T. is funded by a project from the Argentine Fund for Science and Technology (FONCyT), PICT 2014-1818. AA is funded by the Swedish Research Council (B0569601), the European Research Council under the European Union’s Seventh Framework Programme (FP/2007-2013, ERC Grant Agreement n. 331024), the Swedish Foundation for Strategic Research and by a Wallenberg Academy Fellowship. N.S. received funding from the Swiss National Science Foundation (CR32I3-143768).

## SUPPLEMENTARY TEXT

### Bayesian estimation of trait evolution

We developed a Bayesian method to jointly estimate the parameters of a Brownian motion model of trait evolution (BM), i.e. rates and trends, their variation across clades, and the ancestral states at all internal nodes. We used different Markov Chain Monte Carlo (MCMC) algorithms combined into a single MCMC to sample the different parameters: Birth-Death MCMC to estimate the number and placement of shifts in BM parameters, Metropolis-Hastings MCMC to sample rates, trends and the trait value at the root of the tree, and a Gibbs sampler to sample ancestral states at the other nodes. The different algorithms were chosen for their properties to improve the efficiency of the analysis (see also Section *Performance tests*). All three steps are run during a single analysis and our algorithm randomly jumps across them based on user-defined frequencies.

### Birth-Death MCMC to estimate shifts in rates and trends

We implemented a Bayesian algorithm to jointly estimate the number and placement of shifts in the rate and trend parameters of the BM model. Shifts define monophyletic clades within which the rate or trend parameter is treated as independent of the parameters in the other branches of the tree. To infer the number and placement of shifts, we used Birth-Death MCMC (BDMCMC) (Stephens, 2000), an algorithm that has been previously used to estimate rates shifts in other stochastic processes in an evolutionary biology context (Silvestro et al., 2014). Unlike the reversible-jump MCMC (Green, 1995), the BDMCMC-moves across models are not based on an acceptance probability, but on varying rates of a stochastic birthdeath process. The birth rate, determines the probability of proposing a new shift in rates or trends and is fixed to 1 (Stephens, 2000), while individual death rates are calculated for each class of parameters defined by a shift. Death rates determine the probability of removing a rate or trend shift. We use a Poisson distribution with shape parameter set to 1 as prior distribution on the number of rate and trend shifts.

To compute the death rate of a shift is obtained by calculating the likelihood of the tree under a BM with and without the shift. To compute the likelihood without a shift we set the rate (or trend) of the clade identified by the shift to the background rate (or trend), i.e. the current parameter value at its parent node. The death rate of a parameter class is computed as the ratio between the likelihood without the shift and the likelihood with the shift (Stephens, 2000; Silvestro et al., 2014). Thus, rate or trend shifts that improve the fit of the model have a very low extinction rate, and are unlikely to be removed during the BDMCMC. In contrast, rate shifts that do not improve the tree likelihood (or even decrease it) result in high extinction rates and will be removed very quickly by the BDMCMC algorithm.

The algorithm starts with the simplest BM model (i.e. with homogeneous rate and trend parameters) and randomly selects a clade for which a new rate or trend is sampled from their prior distribution. In this case we use an exponential distribution for rates and a normal distribution centered in 0 for trends. The introduction of a shift in the model represents a “birth” event. As soon as there is at least one shift the death rates for each clade identified by a shift are calculated and the following event of the birth-death process will be determined by the relative magnitude of the rates. Additional details about the BDMCMC algorithm are described by (Stephens, 2000; Silvestro et al., 2014).

### Metropolis-Hastings MCMC to estimate BM parameters

For a given set of rate and trend parameters and a vector of ancestral states, the likelihood of a BM model can be calculated as a product of normal densities moving from the tips to the root (*see main text*). We sampled the rate and trend parameters and the ancestral state at the root of the tree using MCMC with acceptance probabilities defined by the posterior odds (Metropolis et al., 1953; Hastings, 1970). We used multiplier proposals for the rate parameters (while properly adjusting the Hastings ratio (Ronquist et al., 2007)) and sliding window proposals for trends and the root state.

### A Gibbs sampler for ancestral states

Sampling the ancestral states from their posterior distribution using the typical acceptance ratio of a Metropolis-Hastings MCMC can be difficult due to the large number of parameters (one for each internal node in the tree), which increase exponentially with the number of tips. Thus, we implemented a Gibbs sampler, in which the ancestral states are sampled directly from their posterior density. This is possible because the posterior probability distribution of an ancestral state under a BM model, given a normal prior distribution, is itself normally distributed. Indeed, because the expected trait value of a BM model after a time *t* is normally distributed (see Eq. 1 in the main text), the posterior density of an ancestral state *x*_*i*_ derives from the combination of four normal distributions: To sample the ancestral states from the posterior we therefore draw random values from the conjugate distribution:

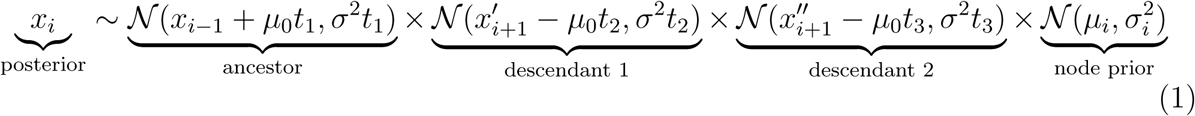

In our implementation, a Gibbs move implies updating all ancestral states iteratively, sampling from Eq. 1.

### Evaluation of the method

#### Simulated data sets

We assessed the accuracy and the efficiency of our Bayesian framework by analyzing simulated data sets and comparing the estimated rates, trends, and ancestral states with the true values. For each simulation, we generated a complete phylogenetic tree (extinct and extant taxa) under a constant rate of birth-death with 100 extant tips (*sim.bd.taxa* function with parameters, *λ* = 0.4, and *μ* = 0.2, in the R package TreePar, (Stadler, 2011)). The number of fossils simulated on the tree was defined by a Poisson distribution with expected number of occurrences (*N*) equal to the total branch length times the preservation rate (*q*). We fixed the number of expected fossils *N* = 20 by setting *q* to the ratio between *N* and the sum of branch lengths. Then, each fossil was placed randomly on the tree and extinct tips without fossils were pruned out (note that this procedure is equicalent to that implemented in the FBD original paper (Heath et al., 2014)). Finally, the fossilized tree was re-scaled to an arbitrary root height of 1.

Phenotypic data were simulated on every tree with the following parameters: rate of evolution (*σ*^2^), phenotypic trend (*μ*_0_), number of shifts in rate, number of shifts in trend, magnitude of the shifts. We simulated data sets under six evolutionary scenarios:

1. constant *σ*^2^ drawn from a gamma distribution Γ(2, 5), and fixed *μ*_0_ = 0
2. fixed *σ*^2^ = 0.1, and constant *μ*_0_ drawn from a normal distribution *𝒩* (2, 0.5)
3. initial 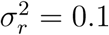, fixed *μ*_0_ = 0, and one rate shift in a randomly selected clade so that 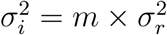, where *m ∼ 𝓊* (8, 16).
4. baseline *σ*^2^ = 0.1, fixed *μ*_0_ = 0, and two shifts in *σ*^2^ drawn from an uniform distribution representing a change with magnitude between 8 and 16 fold
5. fixed *σ*^2^ = 0.1, fixed initial *μ*_0_ = 0, and one shift in *μ*_0_ drawn from a normal distribution *𝒩* (2*r,* 0.5), where *r* was randomly set to −1 or 1 (simulating negative or positive trends, respectively).
6. fixed *σ*^2^ = 0.1, fixed *μ*_0_ = 0, and two shifts in *μ*_0_ each one drawn from a normal distribution *𝒩* (2*r,* 0.5), where *r* was randomly set to −1 or 1 (simulating negative or positive trends, respectively).

For the last two scenarios the location of shifts (in *σ*^2^ or *μ*_0_) was randomly selected by choosing clades that contained between 25 to 50 tips. The branches within the chosen clades were assigned with the new parameter value before performing the phenotypic data simulation using a mapped tree (*make.era.map* function in R package phytools (Revell, 2012)).

We simulated 100 data sets under each of the six scenarios. We additionally ran simulations for scenarios (1) and (2) with a lower number of fossils (N= 5, 1, and 0) adjusting the *q* parameter. For simulations with one fossil, we simulated fossils under a Poisson process with *N* = 20 and kept only the oldest occurrence in the trait analysis.

#### Analysis of simulated data

We analyzed each simulated dataset to estimate the rate and trend parameters of the BM model (*σ*^2^ and *μ*_0_) and the ancestral states. Each dataset was run for 500,000 MCMC generations, sampling every 500 steps. We summarized the results in different ways. First, for each simulation we graphically inspected the results by plotting the phylogeny with the width of the branches proportional to the true and estimated rates. We plotted the true versus estimated *σ*^2^, and *μ*_0_ for each branch on the tree (Fig. S4). The true and estimated ancestral states are compared in a phenogram plot (Revell, 2012).

Secondly, we numerically quantified the overall accuracy of the parameter estimates across all simulations using different summary statistics for each set of parameters. The the BM rate parameters (*σ*^2^) we calculated the mean absolute percentage error (MAPE), defined as:

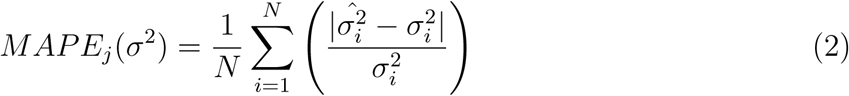

where *j* is the simulation number, 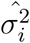 is the estimated rate at branch *i*, 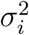 is the true rate at branch *i*, and *N* is the number of branches in the tree. Because the trend parameter can take both negative and positive values, we used the mean absolute error (MAE) to quantify the accuracy of its estimates:

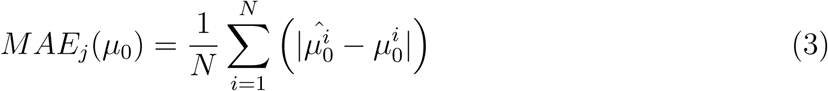

where 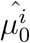 is the estimated trend at branch *i* and 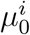 is the true trend at branch *i*. We quantified the accuracy of the ancestral state estimates in terms of coefficient of determination (*R*^2^) between the true and the estimated values. These summary statistics were computed for each simulation scenario (across 100 replicates) and are provided in Figs. S2 and S3.

Finally, we assessed the ability of the BDMCMC algorithm to identify the correct BM model of evolution in terms of number of shifts in rate and trend parameters. We calculated the mean probability estimated for models with different number of rate shifts (*K*_*σ*^*2*^_ ranging from 0 to 4) and shifts in trends (*K*_*μ*0_ ranging from 0 to 4). Note that *K* = 0 indicates a model with constant rate and/or trend parameter across branches. The estimated posterior probability of a given number of rate shifts was obtained from the frequency at which that model was sampled during the MCMC (Stephens, 2000). We averaged these probabilities across 100 simulations under each scenario. We additionally calculated the percentage of simulations in which each model was selected as the best model (i.e., it was sampled most frequently). These summary statistics are provided in the Table S1.

#### Performance tests

We compared the performance of our implementation using Gibbs sampling for ancestral states with that of an implementation using a Metropolis-Hasting algorithm to update the ancestral states, i.e. the method used in other software such as Geiger (Slater et al., 2012; Slater and Harmon, 2013; Pennell et al., 2014) and phytools (Revell, 2012). We implemented the latter option in our code by updating the ancestral states individually using sliding window proposals and acceptance probability based on the posterior ratio. We ran the two implementations on trees with 50, 100, 500, and 1000 tips for 100,000 MCMC iterations, sampling every 50 iterations and using the true parameter values as starting values in the MCMC to avoid burnin. We simulated the data using a constant Brownian rate (*σ*^2^ = 0.1) and no trend (*μ*_0_ = 0) and ran the analyses setting the MCMC to run using the simplest Brownian model (i.e. no rate shifts and no trends). We then calculated for each simulation the run time and the effective sample size (ESS) of the posterior and use them to estimate the run time necessary to reach an ESS = 1000. These performance tests (summarized in Fig. S5a), show that the Gibbs sampler achieved the target ESS of 1000 in a much shorter time than the alternative Metropolis-Hastings algorithm. Importantly, the time required to reach ESS = 1000 scaled linearly with tree size using the Gibbs sampler and exponentially using the Metropolis-Hastings.

The joint estimation of model parameters (rates and trends) and ancestral states allow us to compute the likelihood as a product of normal densities (Fig. S1) while avoiding the use of a variance-covariance (VCV) matrix (commonly used in comparative methods(Felsenstein, 1988; O’Meara, 2012)), which can be very expensive especially for large trees. We ran simulations to assess the performance of the two approaches to calculate the likelihood of the data (product of normal densities against standard calculation using the vcv matrix). We simulated trees of 50, 100, 500, 1000, and 5000 tips under a pure birth process (only extant taxa) and trait data under a simple constant Brownian model (*σ*^2^ = 0.1*, μ*_0_ = 0). On each data set we ran 100,000 MCMC iterations and recorded the run time. The results of these simulations are summarized in Fig. S1b and show that the two approaches to calculate the model likelihood perform similarly for small tree (100 tips). However, the VCV likelihood calculation time scales exponentially with tree size, while our implementation with the product of normal densities scales linearly with tree size. Thus, for instance, our implementation was 1.7 times faster than the alternative with 500 tips and 16.4 times faster with 5000 tips.

## SUPPLEMENTARY TABLES

**Table S1:** Model of testing across simulations. We summarized the mean probability (Pr) estimated for models with different number of rate shifts (*K*_*σ*^*2*^_ ranging from 0 to 4) and shifts in trends (*K*_*μ*0_ ranging from 0 to 4) across 100 simulations under each scenario (see Supplementary Methods). We additionally calculated the percentage of simulations in which each model was selected as the best model (%bm). Values in bold represent the settings used to simulated the data.

| n. shifts | Scenario 1 |  | Scenario 2 |  | Scenario 3 |  | Scenario 4 |  | Scenario 5 |  | Scenario 6 |  |
| --- | --- | --- | --- | --- | --- | --- | --- | --- | --- | --- | --- | --- |
|  | Pr | %bm | Pr | %bm | Pr | %bm | Pr | %bm | Pr | %bm | Pr | %bm |
| $K_{\sigma^2} = 0$ | **0.72** | **97** | **0.81** | **96** | 0.01 | 1 | 0 | 0 | **0.80** | **97** | **0.76** | **92** |
| $K_{\sigma^2} = 1$ | 0.23 | 3 | 0.17 | 3 | **0.73** | **89** | 0.05 | 7 | 0.17 | 2 | 0.19 | 7 |
| $K_{\sigma^2} = 2$ | 0.04 | 0 | 0.02 | 1 | 0.22 | 9 | **0.68** | **91** | 0.03 | 1 | 0.04 | 1 |
| $K_{\sigma^2} = 3$ | 0 | 0 | 0 | 0 | 0.03 | 1 | 0.22 | 2 | 0.01 | 0 | 0 | 0 |
| $K_{\sigma^2} = 4$ | 0 | 0 | 0 | 0 | 0 | 0 | 0.04 | 0 | 0 | 0 | 0 | 0 |
| $K_{\mu 0} = 0$ | **0.59** | **98** | **0.61** | **86** | **0.57** | **92** | **0.57** | **94** | 0.04 | 4 | 0.01 | 1 |
| $K_{\mu 0} = 1$ | 0.31 | 2 | 0.3 | 14 | 0.32 | 7 | 0.32 | 6 | **0.55** | **87** | 0.12 | 14 |
| $K_{\mu 0} = 2$ | 0.08 | 0 | 0.08 | 0 | 0.09 | 1 | 0.09 | 0 | 0.3 | 9 | **0.53** | **79** |
| $K_{\mu 0} = 3$ | 0.01 | 0 | 0.01 | 0 | 0.02 | 0 | 0.02 | 0 | 0.09 | 0 | 0.26 | 5 |
| $K_{\mu 0} = 4$ | 0 | 0 | 0 | 0 | 0 | 0 | 0 | 0 | 0.02 | 0 | 0.07 | 1 |

**Table S2:**
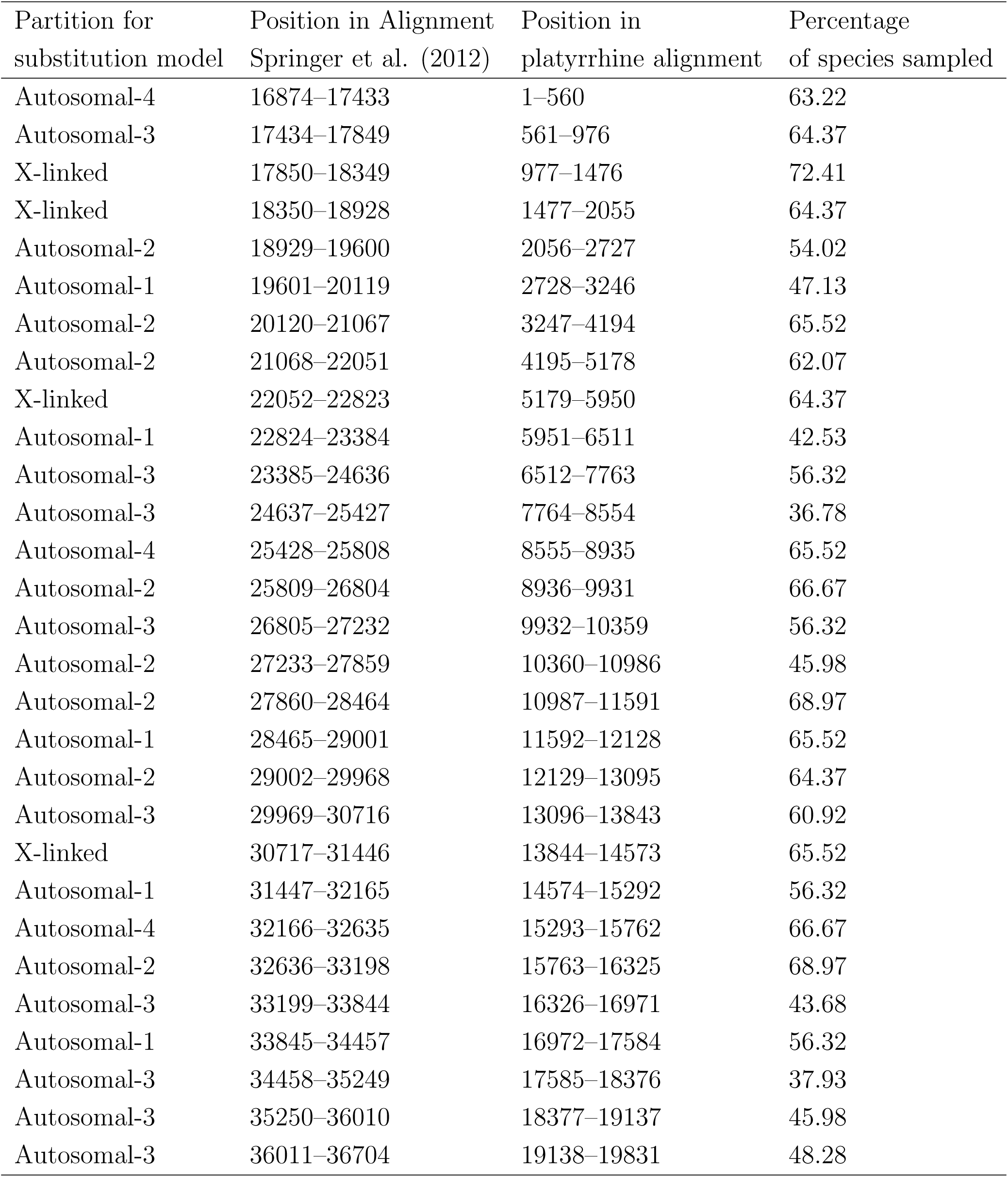
Summary of the molecular data used in this study (Part 1).

**Table S3:** Summary of the molecular data used in this study (Part 2).

| Partition for<br>substitution model | Position in Alignment<br>Springer et al. (2012) | Position in<br>platyrrhine alignment | Percentage<br>of species sampled |
| --- | --- | --- | --- |
| Autosomal-1 | 36705–37359 | 19832–20486 | 67.82 |
| Autosomal-1 | 37360–37916 | 20487–21043 | 59.77 |
| Autosomal-1 | 37917–38456 | 21044–21583 | 68.97 |
| Autosomal-1 | 38457–39061 | 21584–22188 | 68.97 |
| Autosomal-2 | 39062–39711 | 22189–22838 | 68.97 |
| Autosomal-2 | 39712–40049 | 22839–23176 | 63.22 |
| Autosomal-4 | 40050–40362 | 23177–23489 | 62.07 |
| X-linked | 40363–40966 | 23490–24093 | 67.82 |
| Autosomal-3 | 40967–41683 | 24094–24810 | 70.11 |
| Autosomal-2 | 41684–42754 | 24811–25881 | 64.37 |
| Autosomal-2 | 42755–43444 | 25882–26571 | 67.82 |
| Autosomal-2 | 43445–44130 | 26572–27257 | 70.11 |
| Autosomal-1 | 44131–44726 | 27258–27853 | 68.97 |
| Autosomal-1 | 44727–45371 | 27854–28498 | 67.82 |
| X-linked | 45372–45700 | 28499–28827 | 59.77 |
| Y-linked | 45701–46638 | 28828–29765 | 33.33 |
| Y-linked | 46639–47105 | 29766–30232 | 34.48 |
| Autosomal-1 | 47106–47261 | 30233–30388 | 55.17 |
| Autosomal-4 | 47262–48139 | 30389–31266 | 55.17 |
| Autosomal-3 | 48140–48614 | 31267–31741 | 66.67 |
| Autosomal-1 | 48615–49219 | 31742–32346 | 60.92 |
| Y-linked | 49220–49590 | 32347–32717 | 44.83 |
| X-linked | 49591–50401 | 32718–33528 | 67.82 |
| Y-linked | 50402–51254 | 33529–34381 | 39.08 |
| X-linked | 51255–51801 | 34382–34928 | 70.11 |
| mitochondrial | 51802–52938 | 34929–36065 | 82.76 |

**Table S4:**
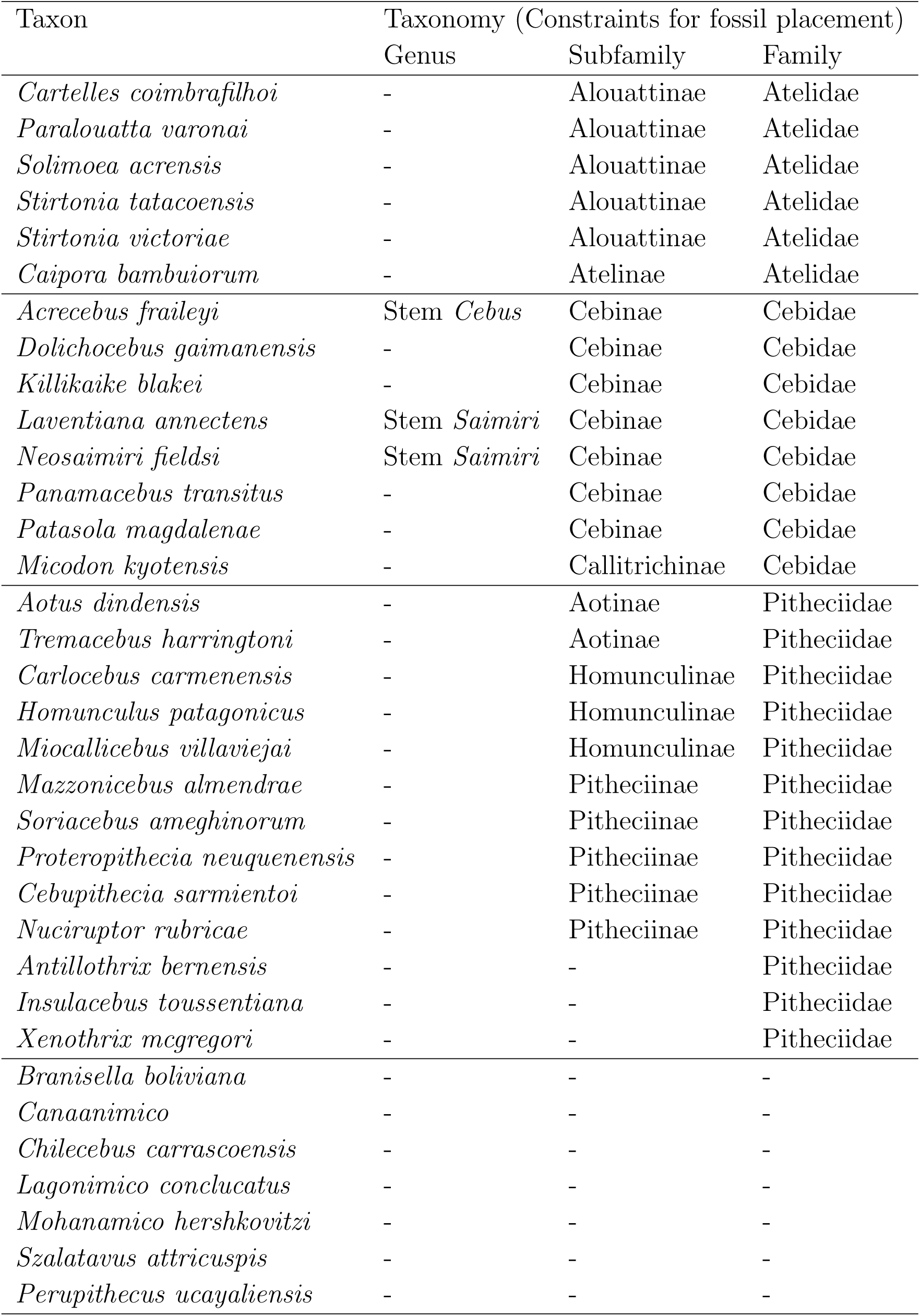
List of fossil taxa used in the phylogenetic analysis of platyrrhines. The taxonomic assignments were used to enofroce topological constraints in the FBD analysis.

**Table S5:**
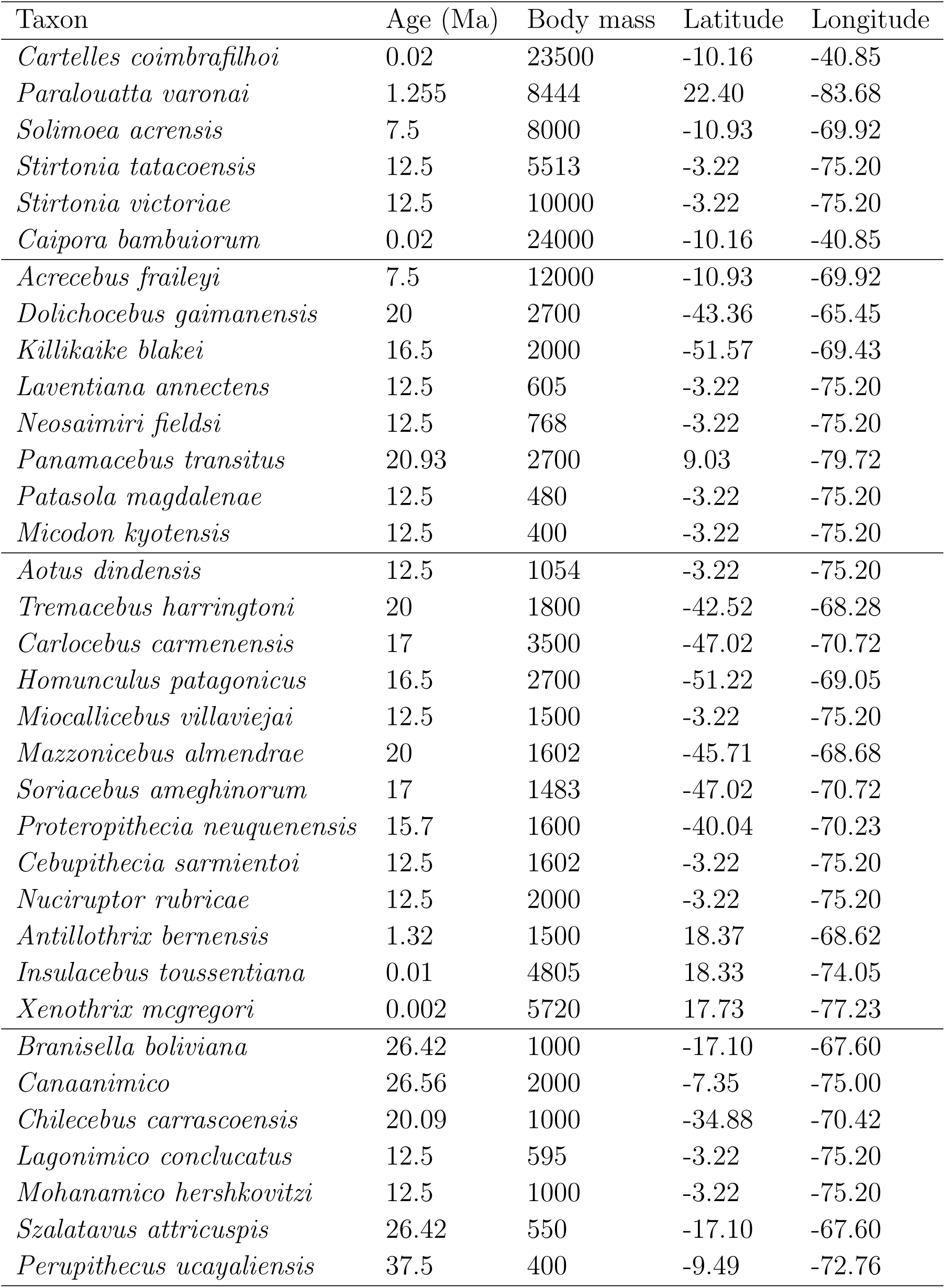
Age, estiamted body mass and geographic coordinates of the fossil records used in phylogenetic and trait evolution analyses.

**Table S6:** Parameters estimated by the fossilized birth-death analysis of extinct and extant platyrrhines. Posterior mean and 95% credible intervals were estimated from the MCMC samples from two independent runs, after removing burn-in.

| Parameter | mean | 95% credible interval |
| --- | --- | --- |
| Net diversification | 0.089 | 0.041 – 0.136 |
| Turnover rate | 0.745 | 0.571 – 0.908 |
| Speciation rate | 0.362 | 0.241 – 0.490 |
| Extinction rate | 0.273 | 0.137 – 0.415 |
| Preservation rate | 0.255 | 0.066 – 0.495 |

**Table S7:**
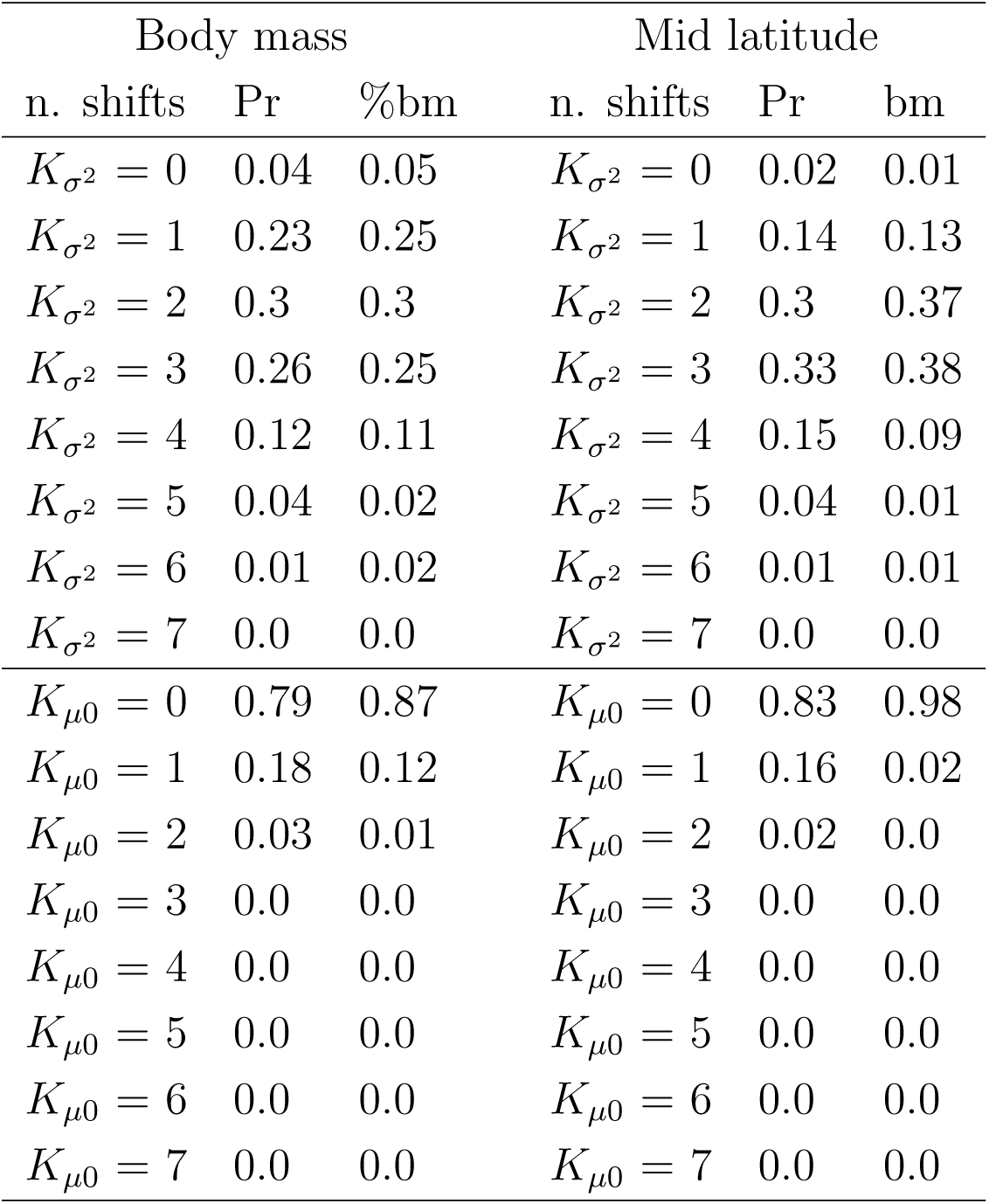
Evolution of body mass and latitude in Platyrrhines: estimated number of shifts in rate and trend parameters averaged over 100 phylogenies of extinct and extant taxa. We summarized the mean probability (Pr) estimated for models with different number of rate shifts (*K*_*σ*^*2*^_ ranging from 0 to 7) and shifts in trends (*K*_*μ*0_ ranging from 0 to 7). For both traits constant rate BM models received very little support and the number of rate shifts ranged between 1 and 4 depending on the tree (Figs. S8, S9). This heterogeneity of BM models estimated across different trees is likely to capture the uncertainties associated with the placement of fossil lineages and branching times. We found little evidence of shifts in trend parameters (S8, S9).

**Table S8:** Posterior estimates of the ancestral body mass for some of the main nodes in the platyrrhine phylogeny. Ancestral states are summarized from 100 analyses as median and 95% credible intervals. For comparison, we show the estimated ancestral states obtained from the analysis of both extinct and extant taxa as well as for analyses based on phylogenies of living taxa only. Note that in trees pruned of all extinct lineages the most recent common ancestor (MRCA) of all extant species coincides with the root node.

| MRCA of | Ancestral body mass (Kg) |  |  |  |
| --- | --- | --- | --- | --- |
|  | using fossils |  | using only extant data |  |
|  | median | 95% CI | median | 95% CI |
| Callicebinae | 1.58 | 0.74 – 3.06 | 1.23 | 0.73–1.86 |
| Pitheciinae | 1.68 | 1.06 – 2.61 | 2.13 | 1.22–3.35 |
| Alouattinae | 6.54 | 3.28 – 10.21 | 6.13 | 4.57–7.86 |
| Cebinae | 2.09 | 0.93 – 3.17 | 1.36 | 0.78–2.12 |
| Aotinae | 0.91 | 0.55 – 1.42 | 0.89 | 0.65–1.16 |
| Callitrichinae | 0.63 | 0.28 – 1.12 | 0.65 | 0.41–0.97 |
| Atelinae | 6.91 | 1.40 – 12.75 | 6.30 | 3.81–9.30 |
| Cebidae | 1.67 | 0.83 – 2.57 | 1.28 | 0.78–1.91 |
| Atelidae | 3.94 | 0.85 – 7.82 | 4.06 | 2.21–6.37 |
| Pitheciidae | 1.59 | 0.58 – 2.69 | 1.70 | 0.93–2.68 |
| All extant species | 1.18 | 0.50 – 2.26 | 1.71 | 0.95–2.71 |
| Clade origin | 0.40 | 0.26 – 0.89 | 1.71 | 0.95–2.71 |

**Table S9:** Posterior estimates of the mid latitude for some of the main nodes in the platyrrhine phylogeny. Ancestral states are summarized from 100 analyses as median and 95% credible intervals. For comparison, we show the estimated ancestral states obtained from the analysis of both extinct and extant taxa as well as for analyses based on phylogenies of living taxa only. Note that in trees pruned of all extinct lineages the most recent common ancestor (MRCA) of all extant species coincides with the root node.

| MRCA of | Ancestral latitude |  |  |  |
| --- | --- | --- | --- | --- |
|  | using fossils |  | using only extant data |  |
|  | median | 95% CI | median | 95% CI |
| Callicebinae | -20.11 | -49.36 – 1.41 | -7.13 | -20.11–6.41 |
| Pitheciinae | -29.42 | -49.90 – -2.52 | -3.33 | -18.84–11.24 |
| Alouattinae | -7.56 | -31.69 – 17.66 | -11.24 | -25.21–7.86 |
| Cebinae | -25.73 | -51.99 – 9.94 | -5.73 | -20.36–8.61 |
| Aotinae | -7.21 | -22.00 – 6.92 | -3.44 | -11.91–5.23 |
| Callitrichinae | -17.78 | -36.07 – -1.20 | -7.39 | -19.65–4.28 |
| Atelinae | -15.42 | -37.13 – 5.62 | -7.04 | -24.65–11.07 |
| Cebidae | -29.40 | -46.99 – -8.87 | -6.08 | -18.66–6.82 |
| Atelidae | -17.60 | -38.49 – 2.34 | -7.53 | -24.08–9.91 |
| Pitheciidae | -34.27 | -53.63 – -12.38 | -5.29 | -20.63–10.43 |
| All extant species | -22.48 | -45.69 – -5.03 | -5.84 | -21.16–9.31 |
| Clade origin | -9.49 | -34.39 – -0.23 | -5.84 | -21.16–9.31 |

**Table S10:** Estimated body mass (g) of Eocene anthropoids and parapithecoids from North Africa.

| Taxon | Epoch | Locality | Body mass (g) | Family |
| --- | --- | --- | --- | --- |
| *Abuqatrania basiodontos* | Late Eoc. | Egypt | 341 | Parapithecidae |
| *Qatrania wingi* | Late Eoc. – Early Olig. | Egypt | 242 | Parapithecidae |
| *Qatrania fleaglei* | Late Eoc. – Early Olig. | Egypt | 510 | Parapithecidae |
| *Biretia piveteaui* | Late Eoc. | Algeria | 383 | *Incertae sedis* |
| *Biretia fayumensis* | Late Eoc. | Egypt | 273 | *Incertae sedis* |
| *Biretia megalopsis* | Late Eoc. | Egypt | 400 | *Incertae sedis* |
| *Arsinoea kallimos* | Late Eoc. | Egypt | 552 | *Incertae sedis* |
| *Proteopithecus sylviae* | Late Eoc. | Egypt | 800 | Proteopithecidae |
| *Serapia eocaena* | Late Eoc. | Egypt | 1029 | Proteopithecidae |
| *Talahpithecus parvus* | Late Eoc. | Libya | 376 | *Incertae sedis* |

**Figure S1:**
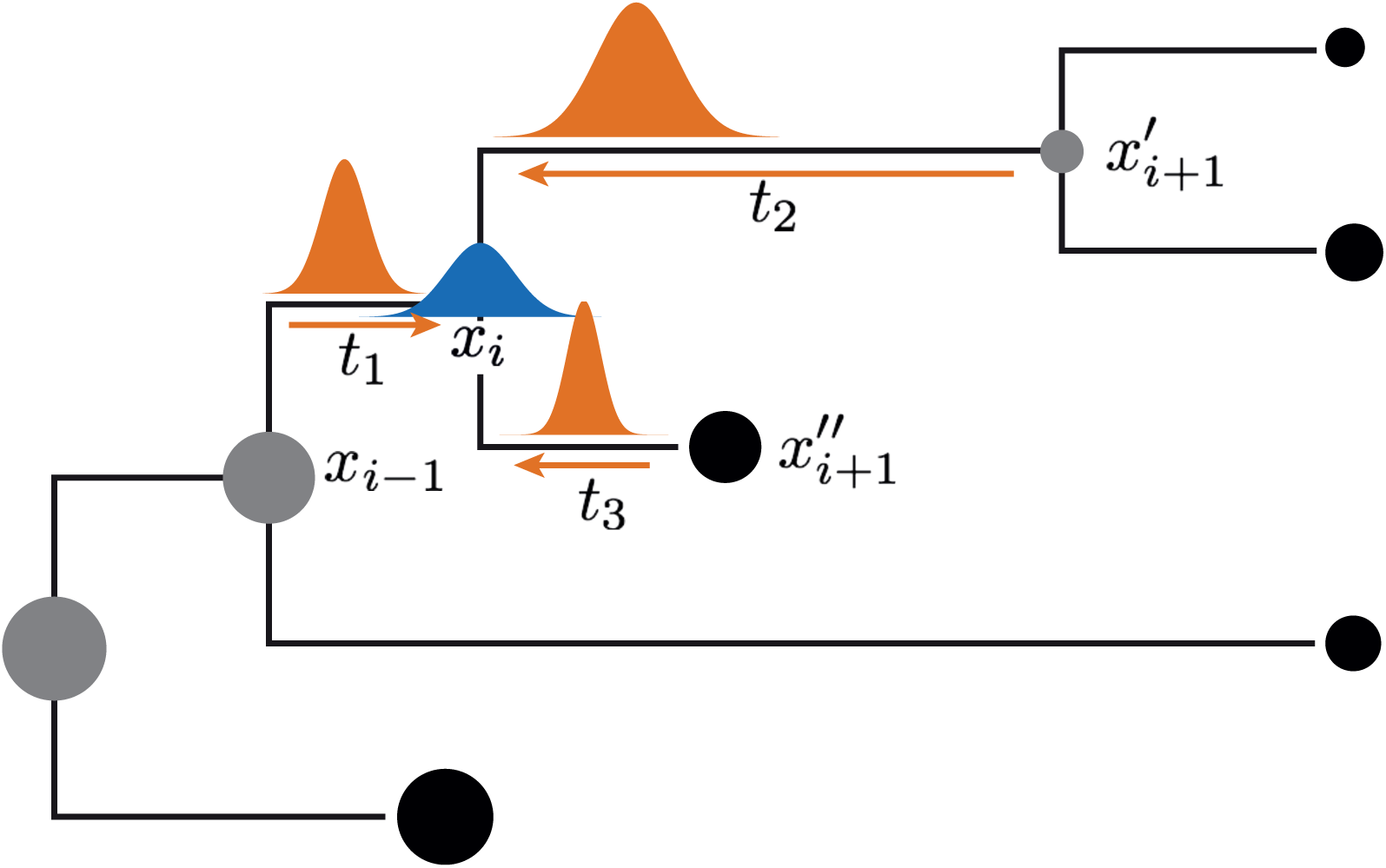
Ancestral states at the internal nodes of the tree are sampled directly from their posterior distribution, which combines four normal densities: two from the descendants, one from the parent node (all of which are based on the current trait states and parameters of the BM model), and one being a vague normal prior distribution on the node state (blue graph). The notation follows that of equation (1).

**Figure S2:**
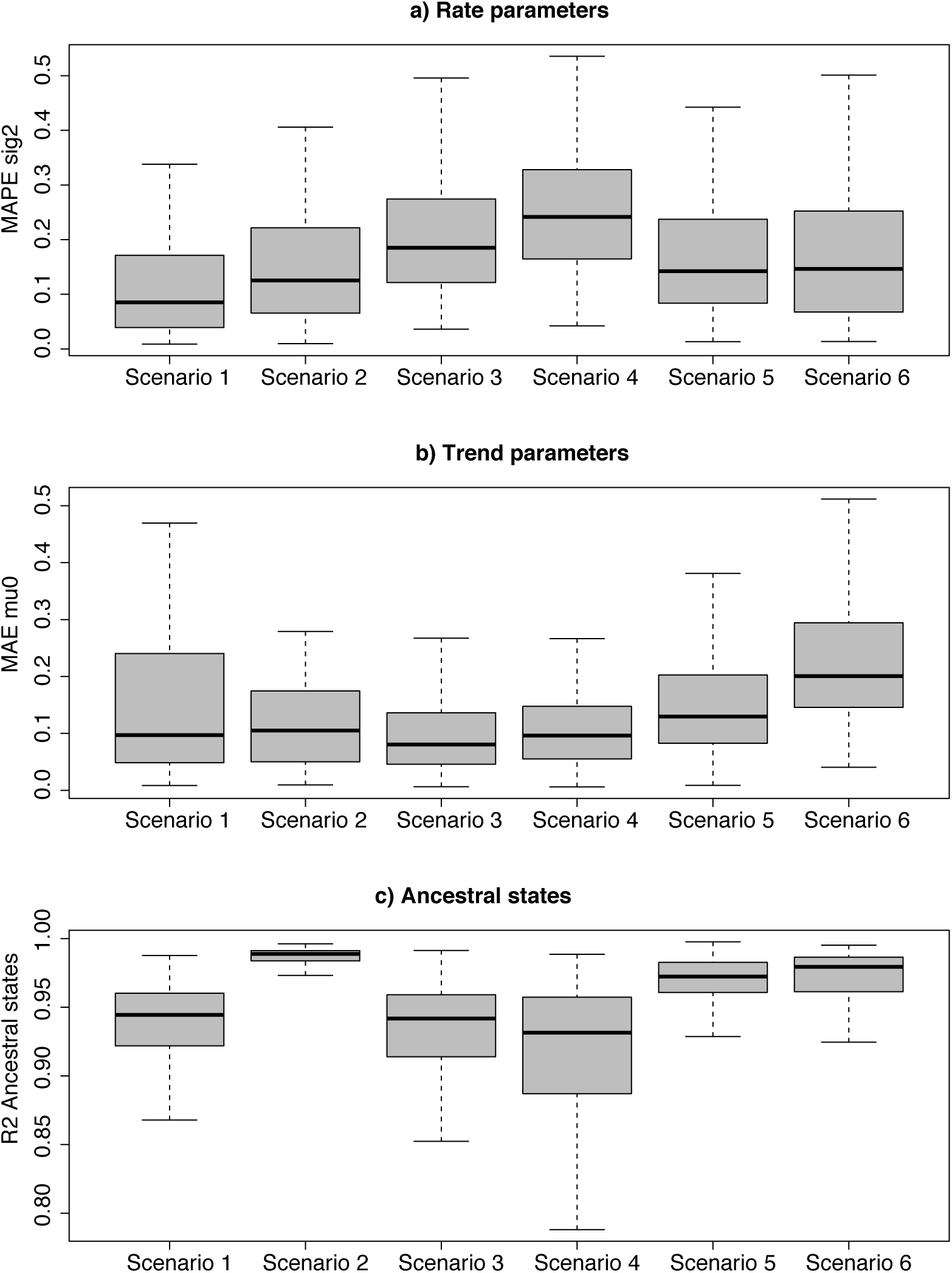
Accuracy of parameter estimation summarized across 100 simulations under six simulation settings (see *Supplementary Methods*). Mean absolute percentage errors (MAPE) are reported for the rate parameters (a; *σ*^2^), Mean absolute errors (MAE) are used for the trend parameters (b; *μ*_0_) and the coefficient of determination *R*^2^ is used for ancestral states (c).

**Figure S3:**
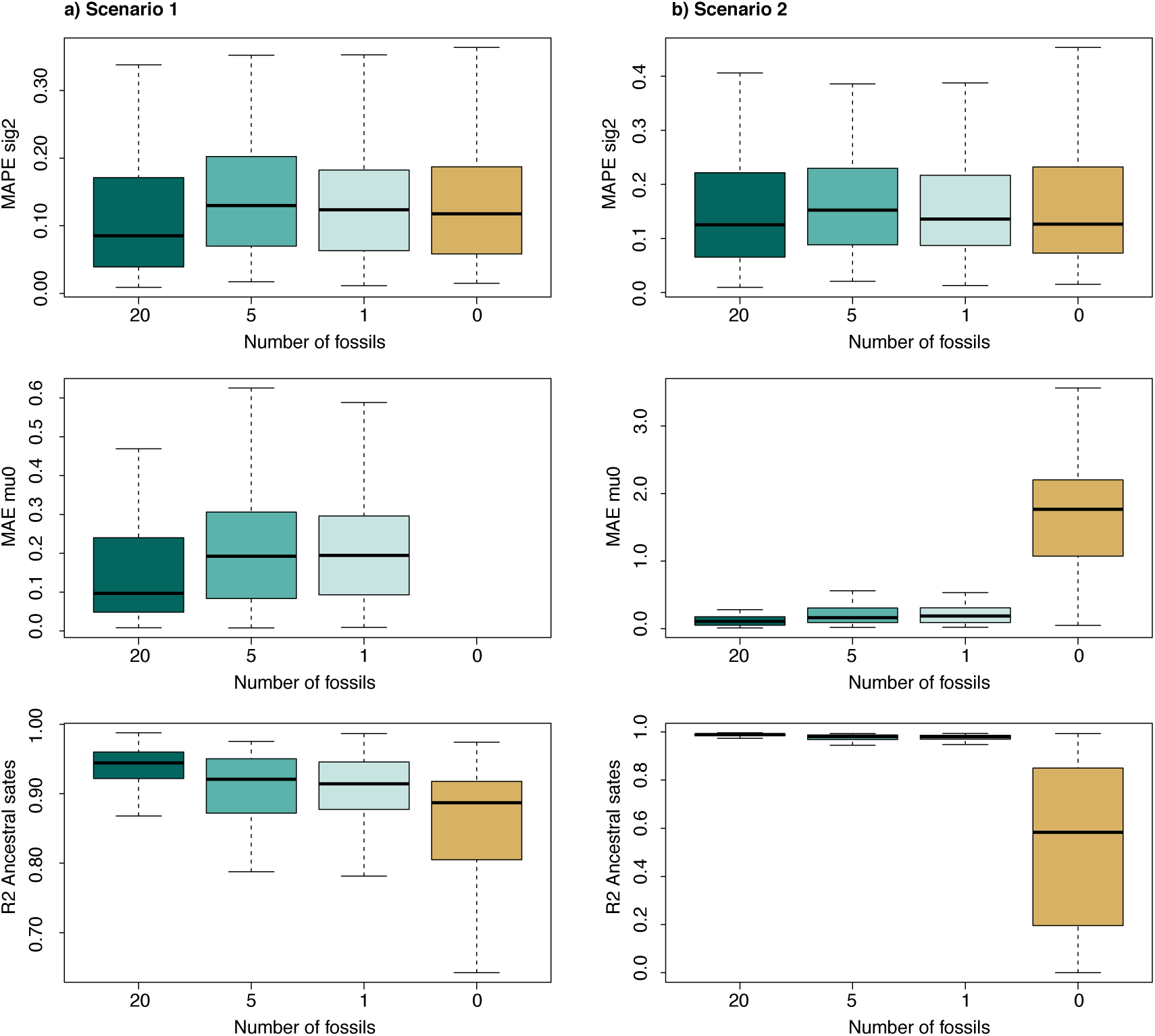
Accuracy of parameter estimation summarized across 100 simulations under scenarios 1 and 2 with decreasing number of fossils: 20, 5, 1, and 0. When the number of fossils was set to 0, only extant taxa were included in the analysis and the trend parameter (*μ*_0_) was not estimated.

**Figure S4:**
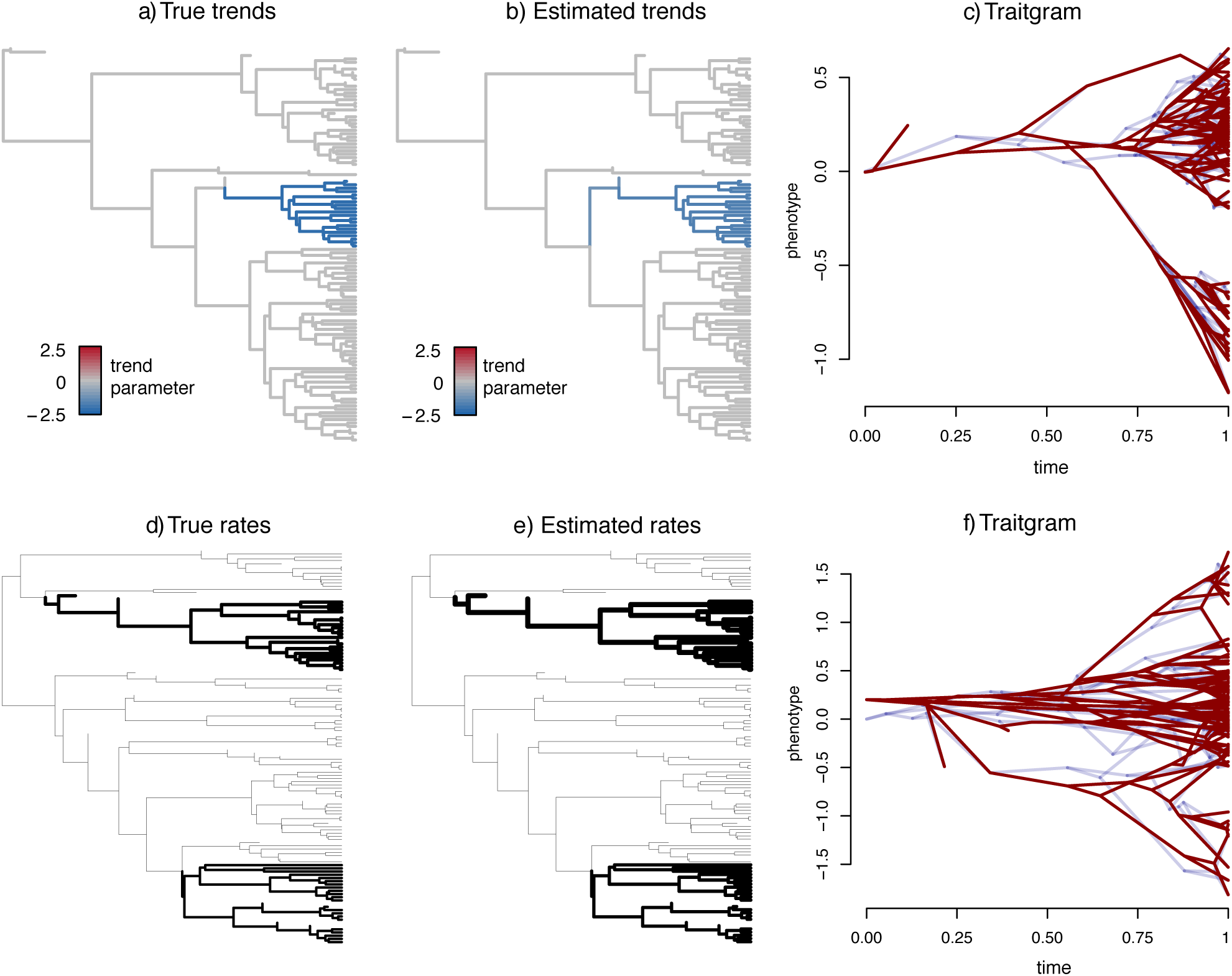
Example of simulated and estimated parameter estimates. Plots in a-c show a simulation from scenario 5, in which one clade evolves under negative trend. Plots in d-f show a simulation from scenario 4, where two rate shifts occur in the phylogeny. The phenogram (c and f) show the true trait evolutionary history(in blue) and the estimated one (in red).

**Figure S5:**
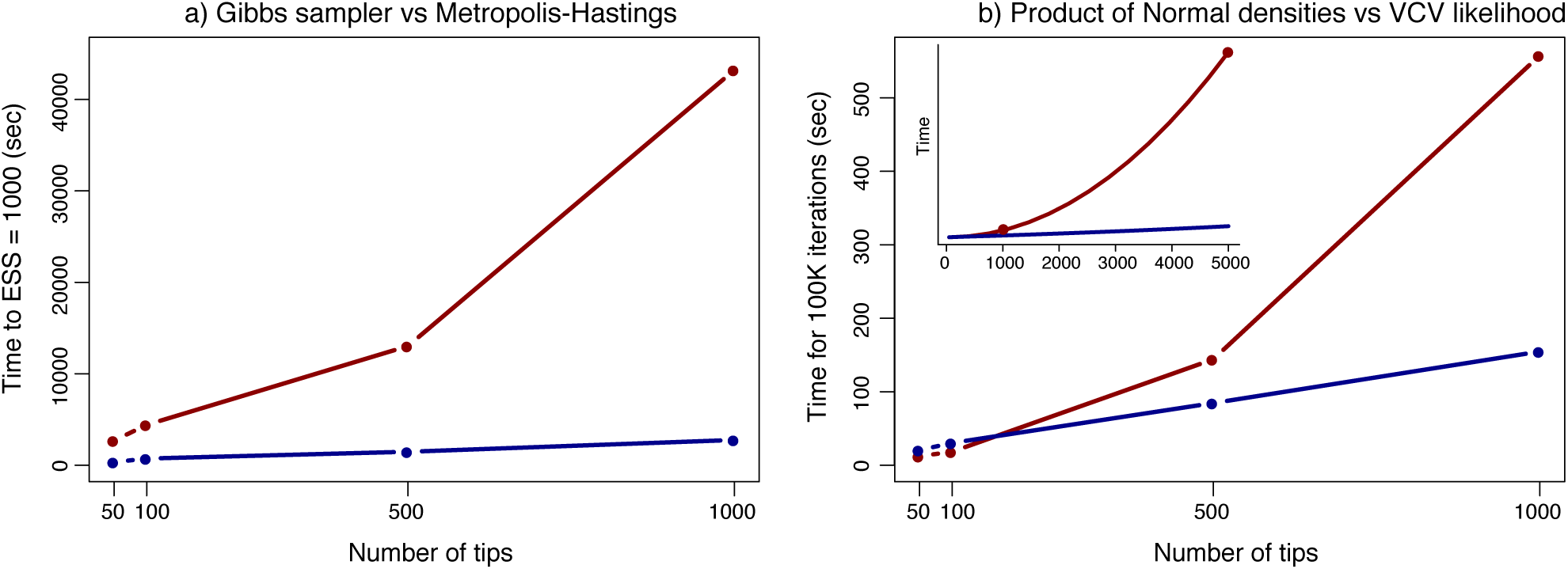
a) Performance of two implementations using Metropolis-Hastings (red) or a Gibbs sampler (blue) to estimate ancestral states

**Figure S6:**
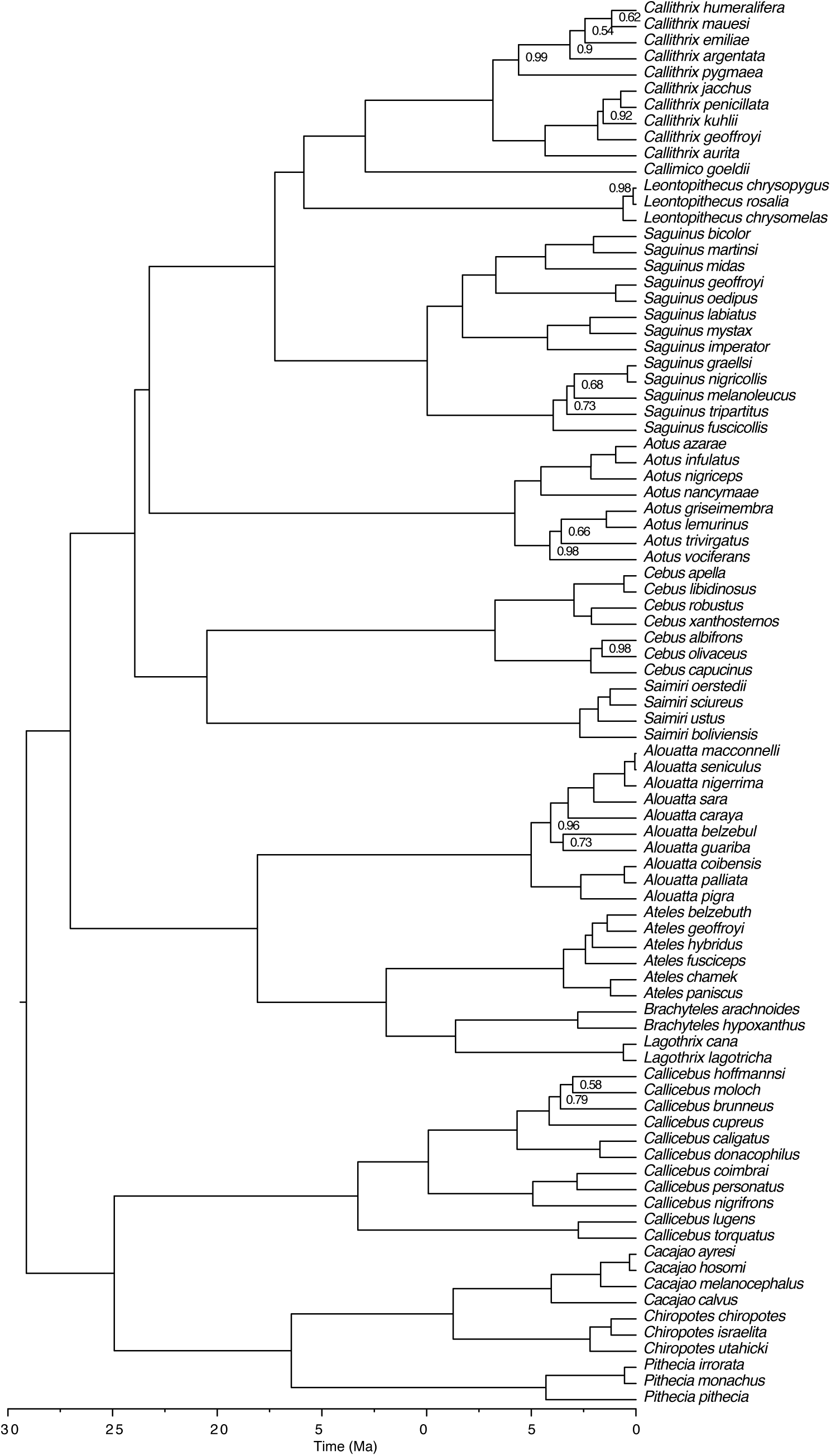
Phylogenetic relationships among extant platyrrhine species. Nodal support (posterior probabilities) is shown only when smaller than 1.

**Figure S7:**
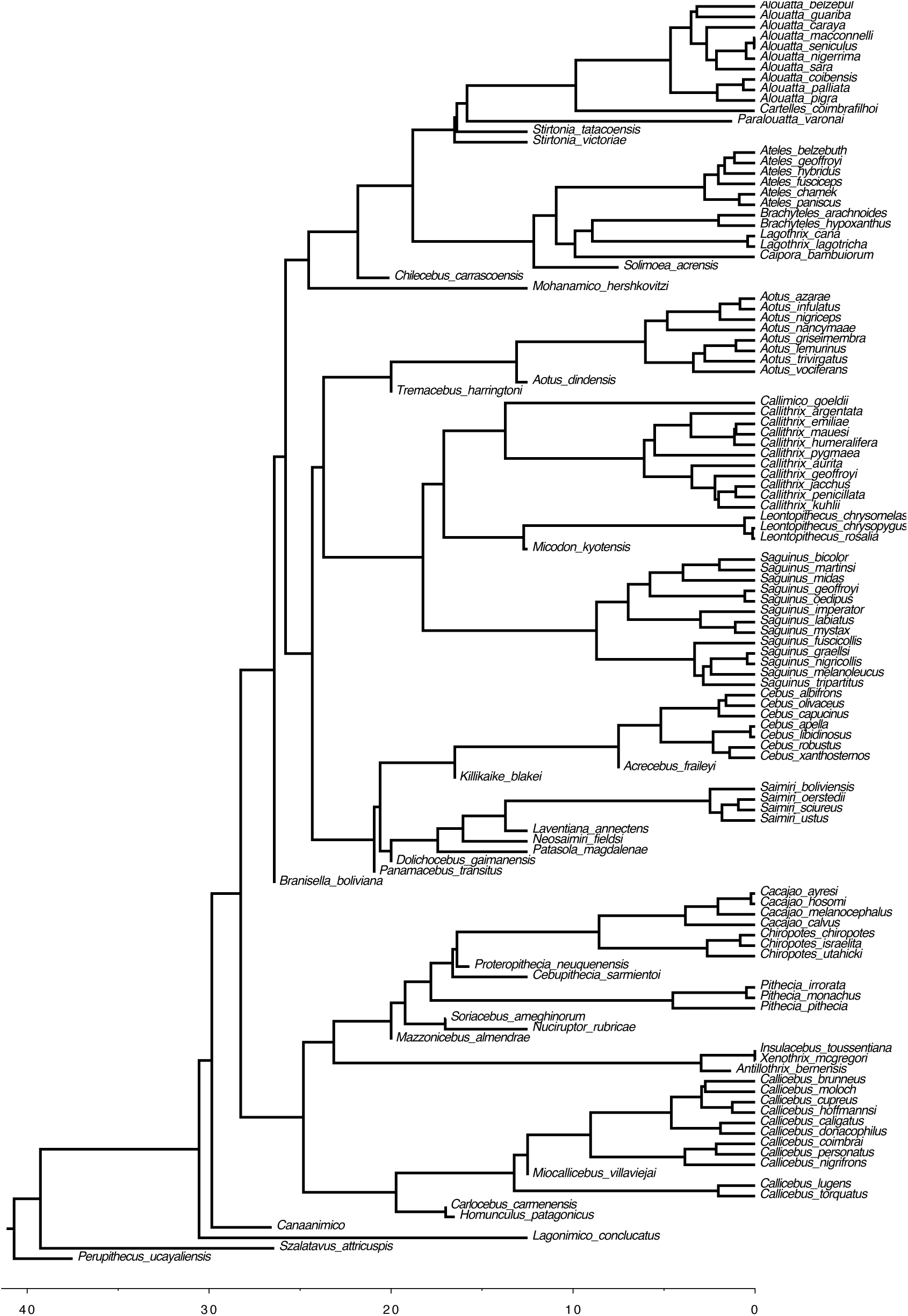
One of the 100 posterior trees of extinct and extant platyrrhines used to perform the trait evolution analyses.

**Figure S8:**
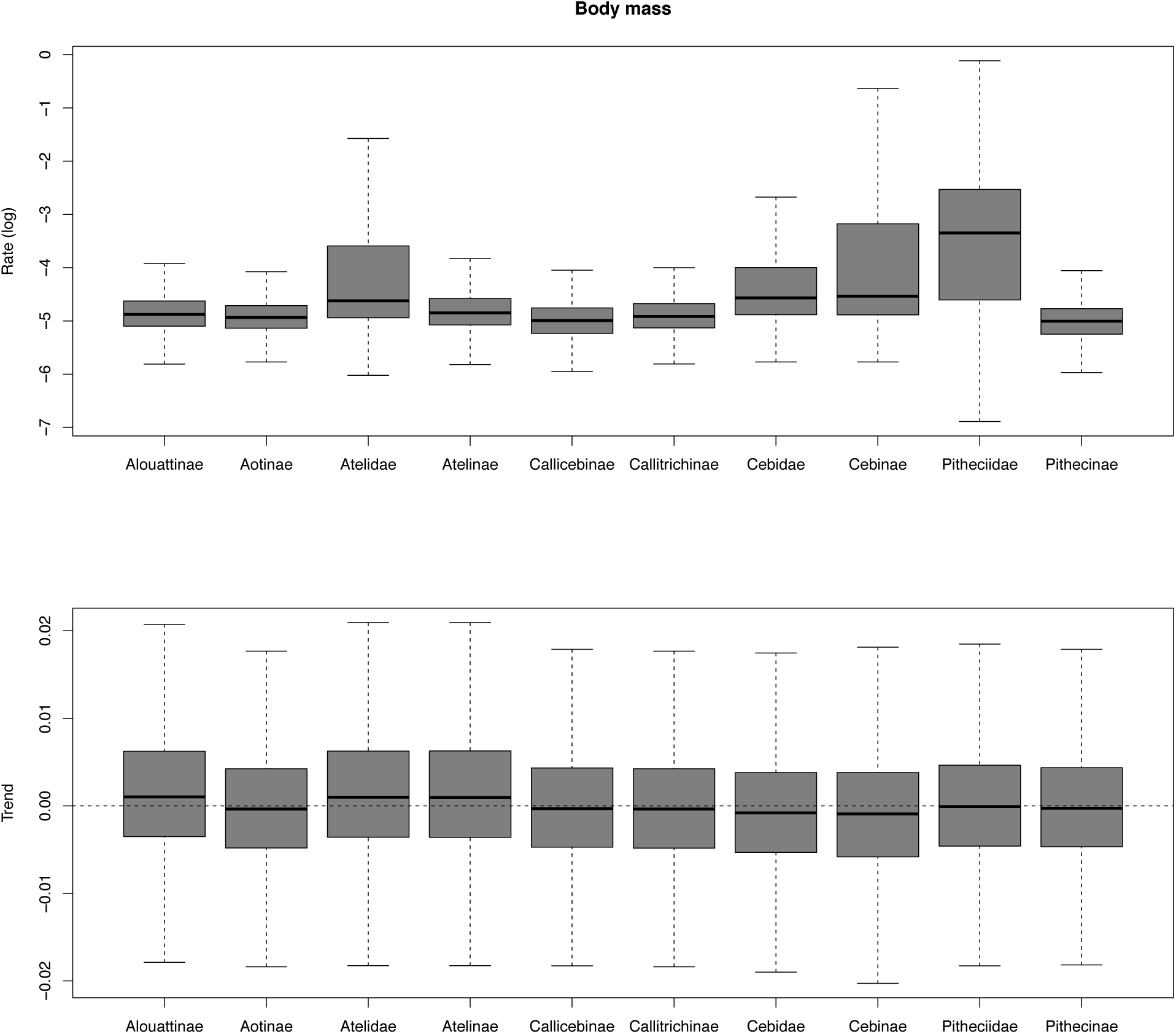
Rates and trend parameters estimated for body mass across families and subfamilies of Platyrrhines, averaged over 100 trees of extant and extinct taxa.

**Figure S9:**
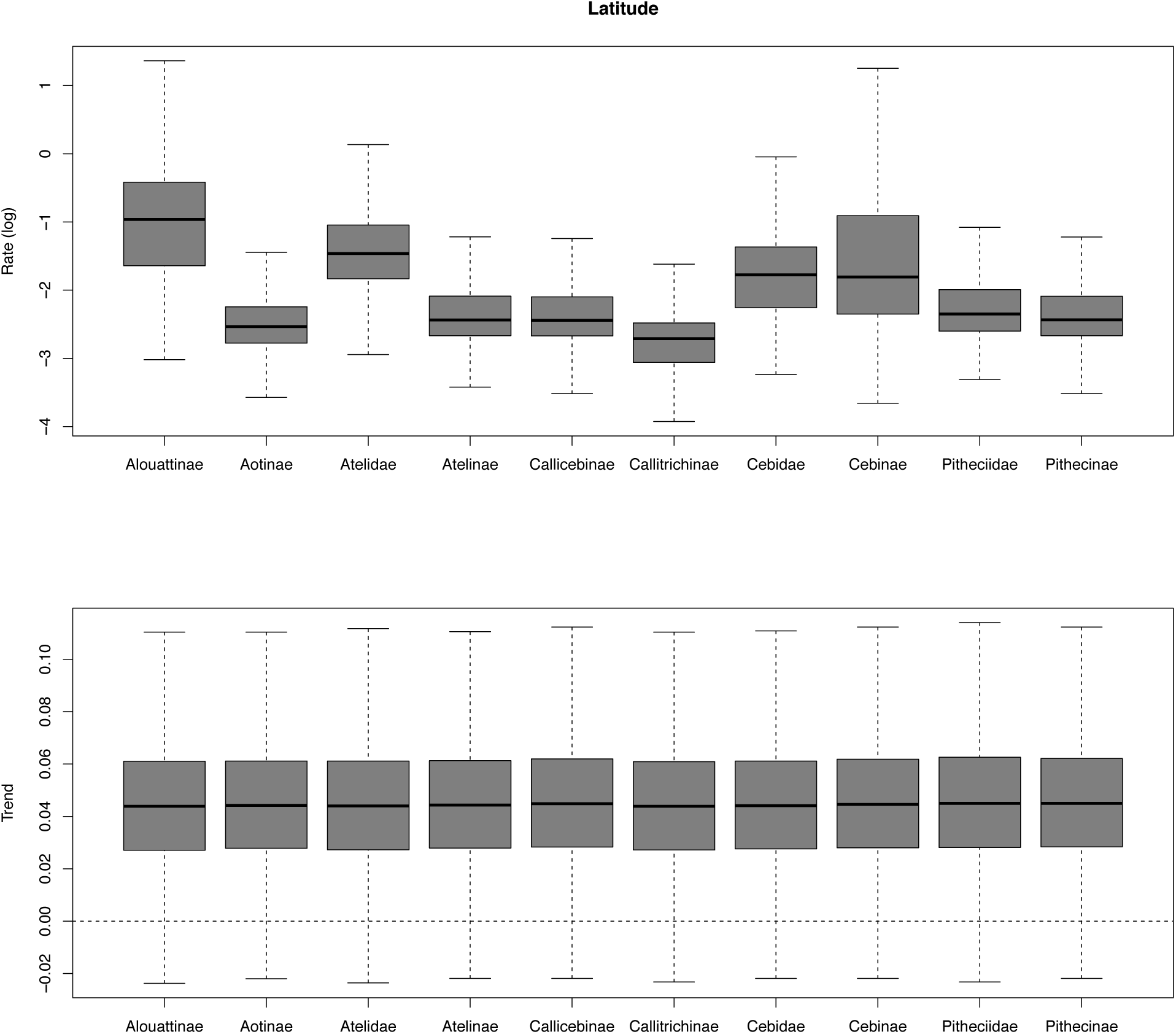
Rates and trend parameters estimated for mid latitude across families and subfamilies of Platyrrhines, averaged over 100 trees of extant and extinct taxa.

**Figure S10:**
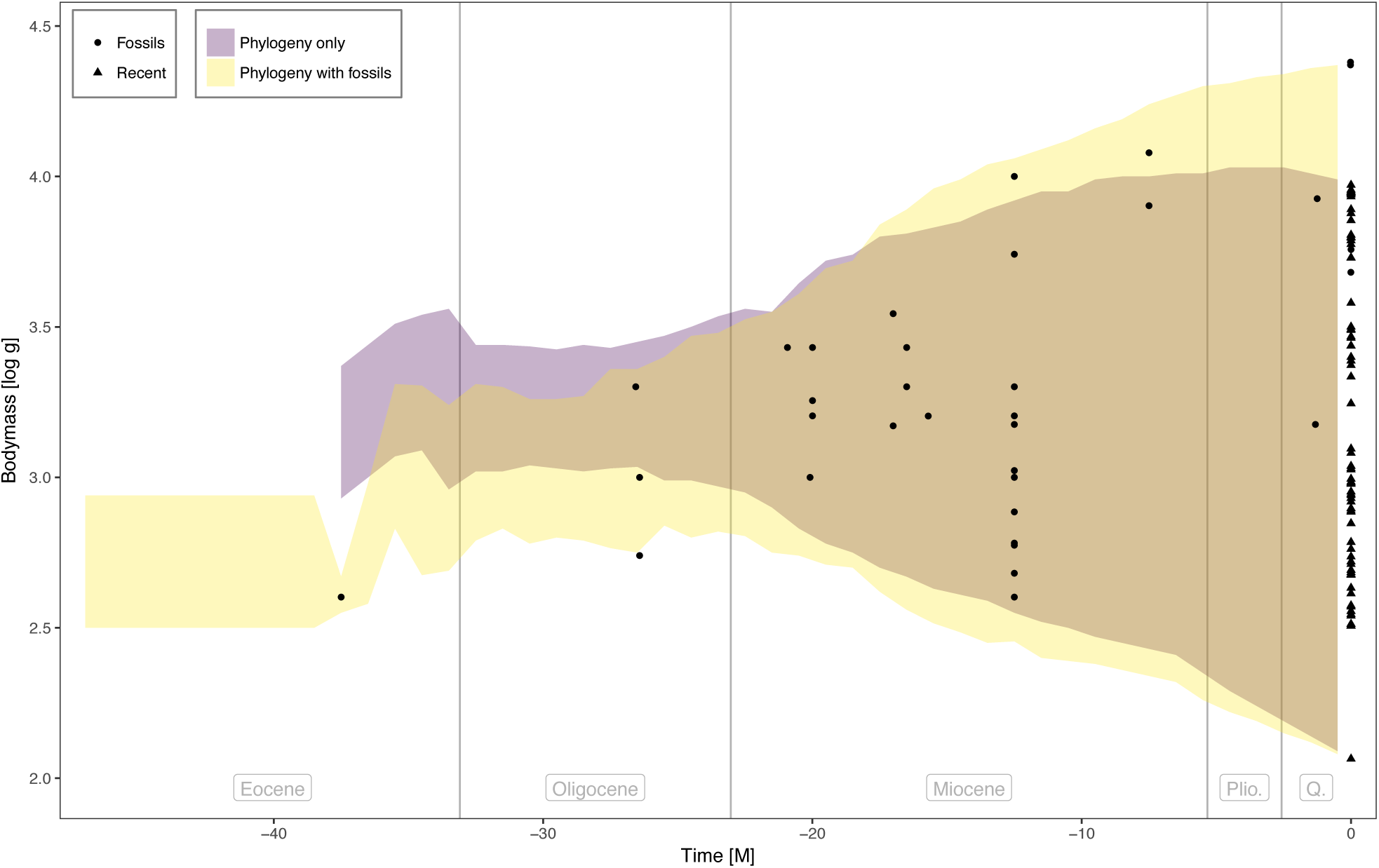
Range of ancestral body mass through time across lineages of platyrrhines. The plot shows a comparison between the estimates obtained when analyzing only extant species and those obtained from the analysis of both extinct and extant taxa. The shaded areas show the minimum and maximum boundaries of the range of estimated body mass averaged over 100 analyses within 1 Myr time bins.

**Figure S11:**
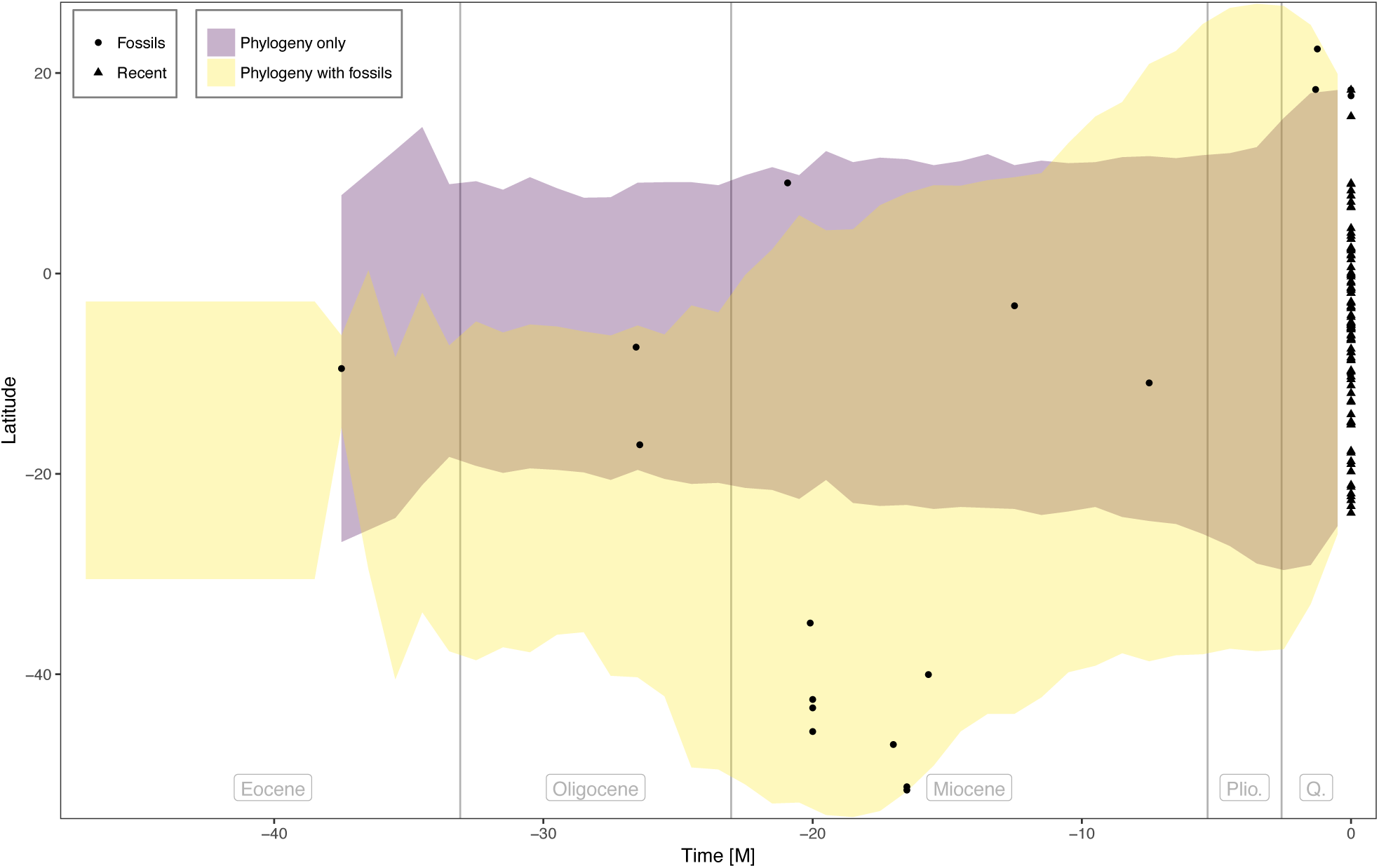
Range of ancestral latitudes through time across lineages of platyrrhines. The plot shows a comparison between the estimates obtained when analyzing only extant species and those obtained from the analysis of both extinct and extant taxa. The shaded areas show the minimum and maximum boundaries of the range of estimated latitudes averaged over 100 analyses within 1 Myr time bins.

